# Activation of vitamin D signaling suppresses platinum-induced ovarian cancer stem cell plasticity

**DOI:** 10.64898/2026.09.22.753468

**Authors:** Tara X. Metcalfe, Sophie Xanders, Renee A. Kinne, Saranya Rajendran, Shu Zhang, Peter C. Hollenhorst, Alexandre Gaspar-Maia, Heather M. O’Hagan, Kenneth P. Nephew

**Affiliations:** Medical Sciences Program, Indiana University School of Medicine-Bloomington, Bloomington, IN 47405, USA; Department of Biology, College of Arts & Sciences, Indiana University, Bloomington, IN 47405, USA; Indiana University Melvin and Bren Simon Comprehensive Cancer Center, Indianapolis, IN 4602, USA; Department of Medical & Molecular Genetics, Indiana University School of Medicine Indianapolis, IN 46202, USA; Department of Anatomy, Cell Biology and Physiology, Indiana University School of Medicine Indianapolis, IN 46202, USA; Division of Experimental Pathology, Department of Lab Medicine and Pathology, Mayo Clinic, Rochester, MN, USA

## Abstract

Advanced-stage high grade serous ovarian cancer (HGSC) initially responds to platinum-based chemotherapy; however, most HGSC tumors recur and develop chemotherapy-resistance, making recurrent ovarian cancer (OC) a fatal disease. Ovarian cancer stem cells (OCSCs), a subpopulation of cells capable of surviving treatment and repopulating tumors, contribute to disease relapse, and understanding the mechanisms(s) responsible for OCSC survival is critical for developing strategies to overcome acquired resistance and prevent tumor recurrence. However, whether platinum-induced OCSC enrichment results from the selection of pre-existing OCSCs or chemotherapy-induced cellular plasticity, and the molecular mechanisms underlying these processes, remain poorly understood. Here, we demonstrated that cisplatin treatment not only enriched for OCSCs but promoted the conversion of ALDH-cells to ALDH+ cells, revealing chemotherapy-induced plasticity as a potential mechanism for OCSC expansion. This conversion was associated with nuclear receptor signaling pathways, vitamin D receptor signaling (VDR/RXR) and retinoic acid receptor signaling (RAR/RXR). Importantly, activation of VDR/RXR signaling with vitamin D reduced stemness phenotypes and the cisplatin induced conversion to ALDH+ cells. In addition, vitamin D combined with cisplatin decreased tumor volume in vivo compared to either treatment alone. In contrast, cisplatin treatment increased RAR/RXR signaling, supporting an associative role for ligand-dependent nuclear receptor signaling in OCSC plasticity. Together, these findings reveal that platinum chemotherapy can actively promote the acquisition of stem-like properties and identifies VDR/RXR signaling as a potential regulatory mechanism of OCSC plasticity. Targeting this pathway with vitamin D may provide a therapeutic strategy to target OCSCs and improve responses to platinum-based chemotherapy.

## Introduction

High grade serous ovarian cancer (HGSC) remains a lethal gynecological malignancy in women^1^. Initially, HGSC is sensitive to platinum-based chemotherapy, but the majority of cases develop resistance over time, which is ultimately fatal^2^. It is widely accepted that in HGSC and other cancers that cancer stem cells (CSC), characterized by their ability to self-renew or differentiate into multiple lineages, contribute to tumor recurrence^3–5^. Characteristics of CSCs include high expression of stemness associated marker genes (e.g. BMI1, NANOG, OCT4) and aldehyde dehydrogenase (ALDH) enzymes^6–8^. We and others have established aldehyde dehydrogenase (ALDH) as a functional stemness marker for the identification and characterization of OCSCs^7, 9–11^. Previous studies have demonstrated platinum treatment increases the percent of ALDH+ cells^8^, and that ALDH+ cells have tumor-initiating activity in vivo^12, 13^. While it has been well-established that platinum drives OCSC enrichment, the underlying mechanisms that mediate the persistence of OCSCs and the emergence of chemo-resistant tumors remain unclear, highlighting the need to develop therapies to prevent recurrence.

The CSC model proposes a unidirectional hierarchy in which CSCs reside at the apex to self-renew and differentiate into non-CSCs in a unidirectional manner^14^. However, recent studies have demonstrated that non-CSCs can acquire stem-like properties through plasticity driven by non-genetic programs, such as, epigenetics and/or transcriptional mechanisms rather than the acquisition of new mutations^15–17^. Mechanisms contributing to CSC plasticity include upregulation of epithelial-to-mesenchymal transition factors, tumor microenvironment influences and chromatin modifications^18–20^. Chemotherapeutic stress has also been observed to promote CSC plasticity, indicating that cancer treatments are capable of reprogramming non-CSCs to be more CSC-like^17, 21^.

Transcriptional programs are important regulators of cell fate, including ligand-dependent nuclear receptor signaling pathways that regulate gene expression^22^. Retinoid X receptor (RXR) functions as a non-permissive heterodimer for multiple nuclear receptors including, retinoic acid receptor (RAR) and vitamin D receptor (VDR)^23, 24^. RAR/RXR signaling is activated by the binding of all-trans retinoic acid (atRA) to RAR and has well-established roles in differentiation, stemness and cell fate decisions^25, 26^. Similarly, ligand-dependent activation of VDR by vitamin D promotes VDR/RXR signaling and regulates differentiation and tumor growth in multiple cancers^27–30^. Despite the established roles of these signaling pathways, the contributions to CSCs and plasticity in response to chemotherapy remains poorly understood.

In the current study, we investigated whether cisplatin treatment could promote OCSC plasticity. By using ALDH-cells isolated from HGSC cell lines, we demonstrate that cisplatin treatment induces the conversion of ALDH- to ALDH+ cells, and that these newly converted cells have increased stemness properties. Furthermore, transcriptomic analysis revealed downregulation of VDR/RXR associated genes, indicating suppression of VDR/RXR was occurring in converted cells, while cisplatin treatment activated RAR/RXR signaling, highlighting distinct ligand-dependent signaling pathways associated with OCSC plasticity. Additionally, we demonstrated that vitamin D in combination with cisplatin inhibited the cisplatin-induced OCSC conversion, reduced stemness and tumor volume in vivo. Together, these data identify nuclear receptor signaling as a potential regulator of OCSC plasticity and support further investigation of vitamin D as a therapeutic strategy to target OCSCs and prevent recurrence.

## Methods

### Cell culture, reagents and drug treatments

HGSC cell lines were obtained through ATCC and maintained at 37°C and 5% CO2 as previously described^31^. OVCAR3 cells were cultured in RPMI media (Thermo Fisher, 11875-119) supplemented with 10% FBS (Gibco, A5670701), 1% antibiotic-antimycotic (Gibco, 15240062) and 2% sodium pyruvate (Gibco, 11360070). OVCAR5 cells were cultured in DMEM media (Thermo Fisher, 11995073) supplemented with 10% FBS (Gibco, A5670701). All cell lines were tested for mycoplasma (Lonza, cat#LT07-318) every 6 months. Stock solutions of cisplatin (MilliporeSigma, 232120) were prepared from 1.67mM stock, diluted with 154mM NaCl and stored at 4°C. HGSC cell lines were treated with IC_50_ doses (OVCAR3: 15μM, OVCAR5: 12μM) for 16 hours^10, 32^. Stock solutions of vitamin D (Enzo Life Sciences, BML-DM200) were prepared in DMSO and cells were treated with IC_50_ doses (OVCAR3: 56nM; OVCAR5: 34nM) for 24 hours. All-trans retinoic acid (Sigma Aldrich, R2625) was prepared in DMSO and cells were treated for 24 hours at varying doses described in figures. Diethlyaminobenzaldehyde (DEAB, MedChemExpress, HY-W016645) was prepared in DMSO and cells were treated for 72 hours. For combination treatments using cisplatin, cisplatin was added the last 16 hours of treatment schemes. For combination treatments of retinoic acid and vitamin D both reagents were added to cells at the same time for 24 hours. For combination treatments of DEAB and retinoic acid, retinoic was added after DEAB treatment. All treatment doses and schemes are detailed in figure legends.

### MTT cell proliferation assay

HGSC ells were seeded onto 96-well plates at a density of 2000 cells/well and treated with increasing concentrations of vitamin D (10-100µM) for 24h, and a 3-(4.5-dimethylthiazol-2-yl)-2.5-diphenyl tetrazolium bromide (MTT; Thermo Fisher Scientific, Waltham, MA, USA, Catalog No. M6494) assay was performed as described previously^33^. The optical density at 450 nm was measured using a BioTek Gen5 plate reader. The IC_50_ values were calculated using GraphPad Prism (v10.6.1).

### Flow cytometry and ALDEFLUOR assay

HGSC cells (1 x10^6^) were seeded in 100mm dishes (Corning, 353003). After treatments, ALDH activity was measured using the ALDEFLUOR assay kit (Stemcell Technologies, 01700) following the manufacturer’s protocol as previously described^8^. ALDH+ cells were analyzed using the LSRII flow cytometer (BD Sciences) or CytoFLEX LX (Beckman-Coulter) and gated based on the DEAB negative control. All flow cytometry data was analyzed using FlowJo software (v10). For cell sorting, 5 x10^6^ cells were used for sorting with the SH800 (Sony). Briefly, HGSC cells were plated in 150mm dishes, and ALDH activity was measured using the ALDEFLUOR assay kit. ALDH+ cells were gated based on the DEAB negative control and collected for functional assays. For non-converted and converted OCSCs, ALDH-cells were plated in 150mm dishes and treated with cisplatin for 16 hours (OVCAR3: IC_50_, 15uM; OVCAR5: IC_50_, 12uM). Following treatment, cells were subjected to the ALDEFLUOR assay kit and sorted using the same gating scheme as before. Cells that had become ALDH+ (converted OCSCs) or remained ALDH-(non-converted) were collected and used for functional assays.

### Annexin V/PI staining

HGSC cells (1x10^6^) were seeded onto 100mm plates (Corning, 353003) and stained with FITC Annexin V Apoptosis Detection Kit with PI (BioLegend, 640914) according to manufacturer’s protocol. Analysis of staining was performed using the CytoFLEX LX (Beckman-Coulter).

### Spheroid formation assay

HGSC cells (2 x 10^3^) cells were pre-treated with cisplatin (OVCAR3: 7.5μM; OVCAR5: 6μM, 3 hours) or vitamin D for 3 hours (OVCAR3: 56nM; OVCAR5: 34nM) and then seeded onto 24-well ultra-low attachment plates (Corning, 3473) in technical triplicates with 1mL of stem cell media as previously described^31^. Stem cell media was supplemented to spheroids every 3 days. At the end of 14 days, spheroids were imaged using the Zeiss Axiovert 40 inverted microscope with Axio-Vision software (Carl Zeiss MicroImaging). Quantification and analysis of spheroids was performed using ImageJ where only spheroids >100μm were counted.

### Migration assay

HGSC cells (5x10^5^) were seeded onto transwell 6.5mm, 8μm pore permeable membrane (Corning, 29442-120) with DMEM complete media (Thermo Fisher, 11995073) containing 10% FBS (Gibco, A5670701) in bottom chamber and starvation media (DMEM only) in top chamber. HGSC cells were incubated for 24h at 37°C. After incubation, membranes were stained using Hema 3 Kit (Fisher Scientific, 122-911) and the percent of migrating cells was normalized to whole cell, untreated samples.

### Colony formation assay

HGSC cells (1.5x10^3^) were seeded onto 6-well plates (Corning, 353046) and allowed to grow for 14 days. At the end of 14 days, cells were washed with PBS, fixed with formalin and stained with crystal violet. Colonies were imaged using SynGene G:Box (GeneSys). Quantification of colonies was performed using ImageJ as previously described^34^.

### RT-qPCR

RNA was isolated from cells using the RNeasy mini kit (Qiagen, 74014) or the RNeasy micro kit (Qiagen, 74004) for sorted cells following the manufacturer’s protocol. Concentration of RNA was determined using the Nanodrop. Total RNA was used for cDNA synthesis using the Maxima First Strand cDNA Synthesis Kit for RT-qPCR (Thermo Scientific, FERK1672). qRT-PCR was performed using Lightcycler 480 SYBR Green I Master kit (Roche Diagnostics, 04707516001) with indicated primers (IDT). Analysis of mRNA expression levels was determined using Lightcycler software version 3.5 (Roche Applied Science) and normalized to *EEF1A1* as previously described^8, 31^. Primer sequences for qPCR can be found in the Supplementary Table S1.

### RNA-seq and data analysis

OVCAR3 cells were sorted using the SH800 (Sony) to obtain ALDH+ ad ALDH-cell populations. Cells were allowed to recover for 3-5 days before ALDH-cells were treated with cisplatin (IC_50_, 16 hours) and sorted to obtain new ALDH+ and ALDH-populations, converted OCSCs and non-converted cells, respectively. All sorted cell populations were collected in biological triplicate, and RNA was isolated using the RNeasy micro kit (Qiagen, 74004) according to the manufacturer’s protocol. RNA-sequencing was performed as previously described^8^. Pathway analysis was performed using Ingenuity Pathway Analysis (IPA) software (Qiagen). Detailed descriptions of bioinformatic analyses can be found in Supplementary Methods.

### Western blot analysis

HGSC cells were treated with cisplatin, all-trans retinoic acid, or vitamin D, lysed in 4% SDS and processed using a QIAshredder (Qiagen, 79656). Protein concentrations were quantified by the DC protein concentration assay (Bio-Rad, 5000113) following the manufacturer’s protocol. Lysates were separated using Mini Protean TGX 4-20% gradient gels (Bio-Rad, 4561094) and transferred to PVDF membrane. Western blots were probed with primary antibodies anti-VDR (Cell Signaling Technologies,12550), anti-RARA (Cell Signaling Technologies, 62294), anti-RXRA (Cell Signaling Technologies, 3085), anti-NCOA2 (Cell Signaling Technologies, 96687) and anti-GAPDH (Cell Signaling Technologies, 5174). After incubation, corresponding HRP-linked rabbit (Cell Signaling Technologies, 7074) and mouse (Cell Signaling Technologies, 7076) secondary antibodies were applied and ECL (Invitrogen, PI32106) and Femto kits (Invitrogen, 34096) were used to visualize bands. Protein bands were quantified with ImageJ and normalized to GAPDH followed by normalization to untreated samples as previously described^31^.

### Proximity Ligation Assay

HGSC cells (2x10^5^) were seeded onto 22mm coverslips (VWR, 10200-036) in 6-well plates (Corning, 353046) and treated with cisplatin, all-trans retinoic acid or vitamin D. After treatments, cells were washed with PBS and fixed for 15 min using 4% paraformaldehyde. Following fixation cells were incubated with methanol for 10 min at -20°C and washed with PBS. Cells were then incubated with 0.5% triton-X for 10 min at RT. The Duolink proximity ligation assay (PLA) was then performed according to manufacturer’s protocol (Sigma Aldrich, DUO92008). Coverslips were incubated with primary antibodies anti-RARA (SantaCruz Biotechnology, sc-515796), anti-RXRA (Proteintech, 21218-1-AP), anti-VDR (SantaCruz Biotechnology, sc-13133) and anti-NCOA2 (Proteintech, 30962-1-AP). PLA signals were imaged with the confocal Leica SP8 microscope using the 63x oil lens and quantified using FIJI (ver 2.16.0). PLA signals were quantified as PLA/puncta and normalized to 500 cells.

### ELISA

HGSC cells were seeded onto 100mm plates (Corning, 353003) and treated with cisplatin or DEAB. After treatments, cells were washed and scraped with PBS into Eppendorf tubes. Samples were then placed in the Bioruptor pico (Diagenode) for 3 cycles 30 sec ON/OFF at 4°C and then centrifuged at 10,000xg at 4°C for 20 min The supernatant was collected and used for the human retinoic acid ELISA kit (CUSABIO, CSB-E167-12h) following the manufacturer’s protocol. Briefly, 50uL of supernatant was mixed with 1X HRP-conjugate in the provided 96-well plate and incubated at 37°C for 60 min. Following incubation, the plate was washed with the wash buffer provided and 3,3’,5,5’-tetramethylbenzidine (TMB) substrate was added to each well. The plate was then incubated at 37°C for 20 min after which a stop solution was added to each well. The absorbance was read using a microplate reader (Biotek) at 450nm and 570nm. Wavelength correction was performed by subtracting the 570nm reading from the 450nm reading. To determine the concentration of retinoic acid in samples a standard curve was generated using serial dilutions of the provided standard in the kit and plotted as OD vs. concentration. The standard curve was fit using an exponential regression model (y=ae^bx^). Curve fitting was performed using Excel and sample concentrations were calculated by rearranging the exponential equation to solve for *x*.

### Luciferase

The RARE sequence was cloned into the firefly luciferase reporter pGL4.25 (Promega) cut with XhoI. The RARE sequence was PCR cloned from pGL3 RARE-RFP (Addgene #183054), and the final plasmid was generated using standard Gibson Assembly protocols using Gibson Assembly Master Mix (NEB). HGSC cells were seeded onto 6-well plates at 2.5x10^5^ cells/well and co-transfected with the RARE pGL4.25 and renilla control vectors pGL4.74[hRluc/TK] using lipofectamine 3000 (ThermoFisher, L3000015) for 24 hours, prior to cell treatments. After treatments, cells were harvested and Dual Luciferase Reporter Assay System (Promega, E1910) was performed as described previously^35^. Luciferase values were analyzed with plate reader (Agilent BioTek) and luciferase values were normalized to renilla values.

### Mouse study

Animal studies were performed under protocol #25037 and all mouse experiments were performed according to ethical guidelines approved by the Institutional Animal Care and Use Committee of Indiana University (Indianapolis, IN, USA). OVCAR3 cells (5x10^6^) were mixed with Matrigel in 1:1 ratio and injected subcutaneously 6–8-week-old female NSG mice. Tumor growth was monitored by caliper measurements of length and width, and tumor volume was calculated using the formula: Volume = L x W x 0.5. Once tumors reached 100mm^3^, mice were randomized into 4 treatment groups, (1) vehicle (sesame oil), (2) cisplatin (2 mg/kg, i.p) once weekly for 3 weeks, (3) vitamin D (0.5μg/kg, i.p) three times per week for 3 weeks and (4) combination of cisplatin + vitamin D. Tumor volumes and body weights were monitored twice weekly until tumors in the control group reach 500 mm^3, and then measured daily (Supplemental Fig. S5). At the end of the study all mice were sacrificed, and tissues were collected for IHC and qRT-PCR analyses as previously described^8, 31^. Survival time was recorded from treatment initiation until humane endpoint.

### Immunohistochemistry

Tumors excised from mice were fixed overnight in 10% formalin, embedded in paraffin, and sectioned by the Histology Lab Service Core facility at Indiana University (Indianapolis, IN, USA). Slides stained with H&E served as controls. Briefly, sections were deparaffinized and antigen retrieval in 10mM sodium citrate (Vector Laboratories, H-3300) was performed in a pressure cooker before the sections were blocked with 5% goat serum with 1%BSA in TBS plus 0.025% TritonX-100. Slides were incubated overnight at 4°C with antibodies against ALDH1A1 (CST, 54135), RARA (ProteinTech, 10331-1-AP), RXRA (Proteintech, 21218-1-AP) and VDR (CST, 12550). Primary antibodies were detected using SignalStain^®^ Boost Detection Reagent (CST, Rabbit: 8114) and developed with SignalStain^®^ DAB Substrate Kit (CST, 8059) followed by dehydration with increasing alcohol solutions and mounted. Slides were imaged with Motic EasyScan Pro scanner and analyzed with QuPath software.

## Statistical analysis

All data are presented as mean values ± SEM of at least three biological experiments unless otherwise indicated. For comparisons between two groups, an unpaired Student’s *t*-test was used. For comparisons of three or more groups, a one-way ANOVA followed by Tukey’s multiple comparisons test in GraphPad Prism (v10.6.1). For all figures, * p<0.05, ** p<0.01, *** p<0.001, **** p<0.0001. All significant comparisons are shown. Statistical details for each experiment are included in the figures and figure legends.

## Data availability

RNA-seq results are available for download at Gene Expression Omnibus (GEO) data repository at the National Center for Biotechnology Information (NCBI) under the accession number GSE336228

## Results

### Platinum induces OCSC plasticity and functional stemness

To test whether cisplatin (CDDP) treatment induces the conversion of ALDH-cells to ALDH+ cells, ALDH-cells were FACS isolated from HGSC cells lines OVCAR3 and OVCAR5 using the ALDEFLUOR assay and placed into culture. In the absence of treatment, ALDH-cells remained ALDH-for up to 10 days, with less than 1% converting to ALDH+, similar to DEAB negative controls (Fig. 1A; Supplementary Fig. S1A; Supplementary Fig. S2 for flow cytometry gating strategy). To assess the effect of CDDP on phenotypic conversion, sorted ALDH-cells were treated with CDDP for 16h at the IC_50_ dose (OVCAR3, 15µM, OVCAR5,12µM) and the ALDELFUOR assay was performed to determine the percent of cells that had converted to ALDH+. Treatment resulted in 10% conversion in OVCAR3 cells and 3-4% conversion in OVCAR5 cells (Fig. 1B; Supplementary Fig. S1B). Cells that remained ALDH-after CDDP treatment were defined as non-converted and cells that had converted to ALDH+ were defined as converted OCSC (Fig. 1C). Annexin V/Propidium Iodide staining confirmed that CDDP had minimal effects on the viability of ALDH-cells at this time point, indicating that a viable population of cells was present for conversion (Supplementary Fig. S3).

**Figure 1.**
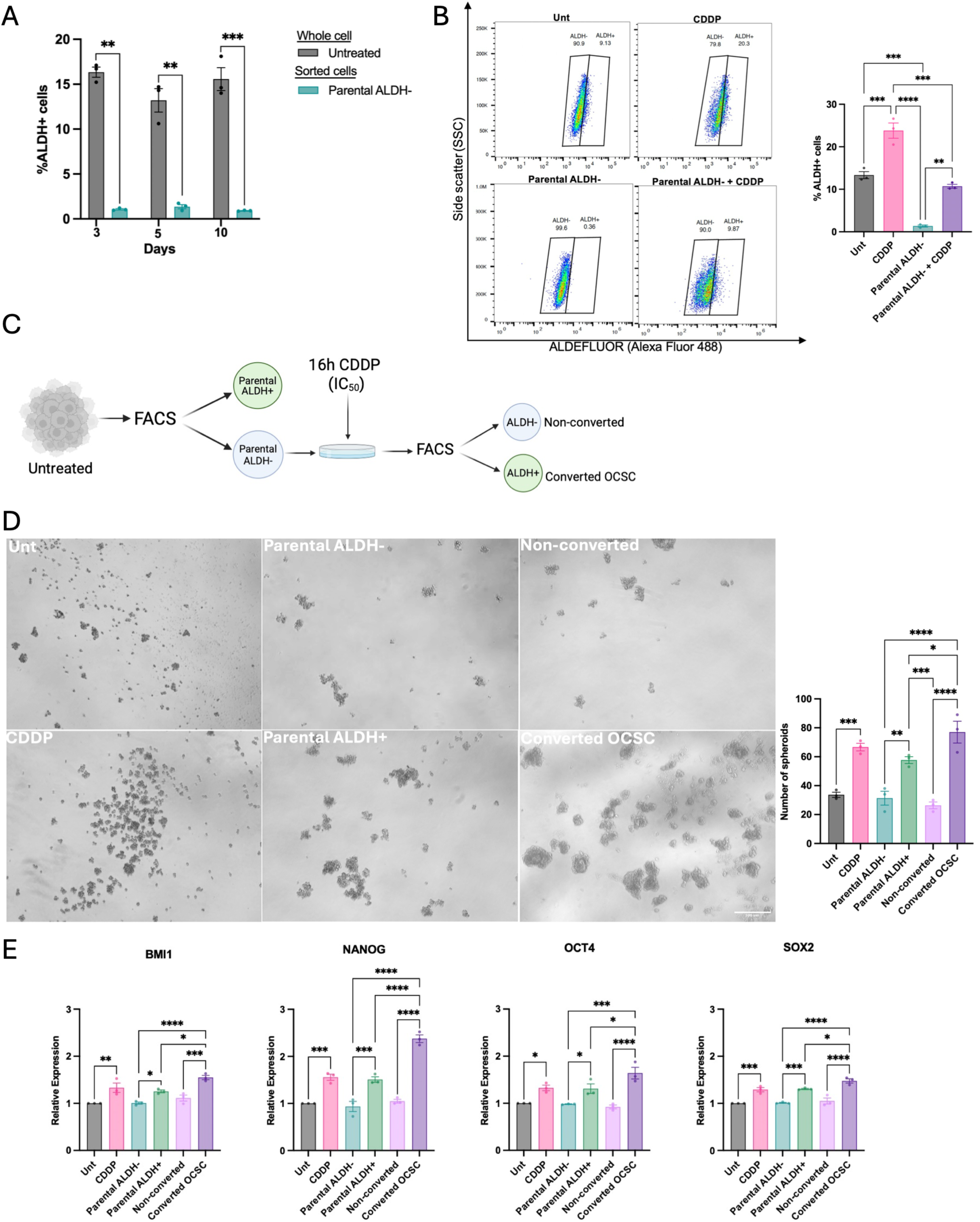
A. OVCAR3 ALDH-cells were FACS isolated and cultured at the indicated timepoints. The percent of ALDH+ cells was determined using the ALDEFLUOR assay compared to an untreated, whole cell population. **B.** The percentage of cells that converted to ALDH+ was determined using the ALDEFLUOR assay compared to untreated and cisplatin (16h, 15µM) treated whole cell populations. Gating strategy: left. **C.** Treatment schematic for FACS isolation of non-converted and converted OCSCs. **D.** OVCAR3 cells treated with CDDP (3h, 7.5µM), vitamin D (3h, IC50, 56nM) or combination and 2000 cells/well were seeded in low adhesion conditions after treatment. Representative images of spheroid formation and quantification (>100um) (right). Scale bar, 500µm. **E.** Expression of stemness associated genes was determined by RT-qPCR analysis in untreated and CDDP treated (16h, 15µM) whole cell populations, parental ALDH+, parental ALDH-, non-converted and converted OCSCs. All experiments were performed in biological triplicate unless stated otherwise, +/- SEM, P *<0.5, P **<0.01, P ***<0.0001, P ****<0.00001.

To evaluate stemness properties of sorted cells, spheroid formation was performed using parental ALDH+, parental ALDH-, non-converted and converted OCSC cell populations (Fig. 1C). Converted OCSCs had a significant increase in the number of spheroids formed compared to sorted cell populations, including parental ALDH+ cells (Fig. 1D; Supplementary Fig. S1D). Similarly, expression of stemness associated genes (BMI1, NANOG ,OCT4, SOX2) in OVCAR3 was increased in converted OCSCs relative to parental ALDH+ cells, suggesting there were distinct differences between OCSC populations (Fig. 1E; Supplementary Fig. S1E).

### Transcriptomic profiling distinguishes converted OCSCs from non-converted populations

To investigate the underlying mechanisms involved in OCSC plasticity, RNA-seq was performed on parental ALDH+, parental ALDH-, non-converted and converted OCSCs. Principal component analysis distinguished converted OCSCs from other populations (Fig. 2A). Ingenuity Pathway Analysis software was used to identify differentially expressed genes and pathways associated with each cell population. Pairwise comparisons were performed between of non-converted vs parental ALDH+ and converted vs parental ALDH+ populations, using parental ALDH+ cells as a baseline control to distinguish genes associated with pre-existing stem-like features. Overall, non-converted cells had 848 uniquely expressed genes, whereas converted OCSCs had 1,205 uniquely expressed genes (Fig. 2B).

**Figure 2.**
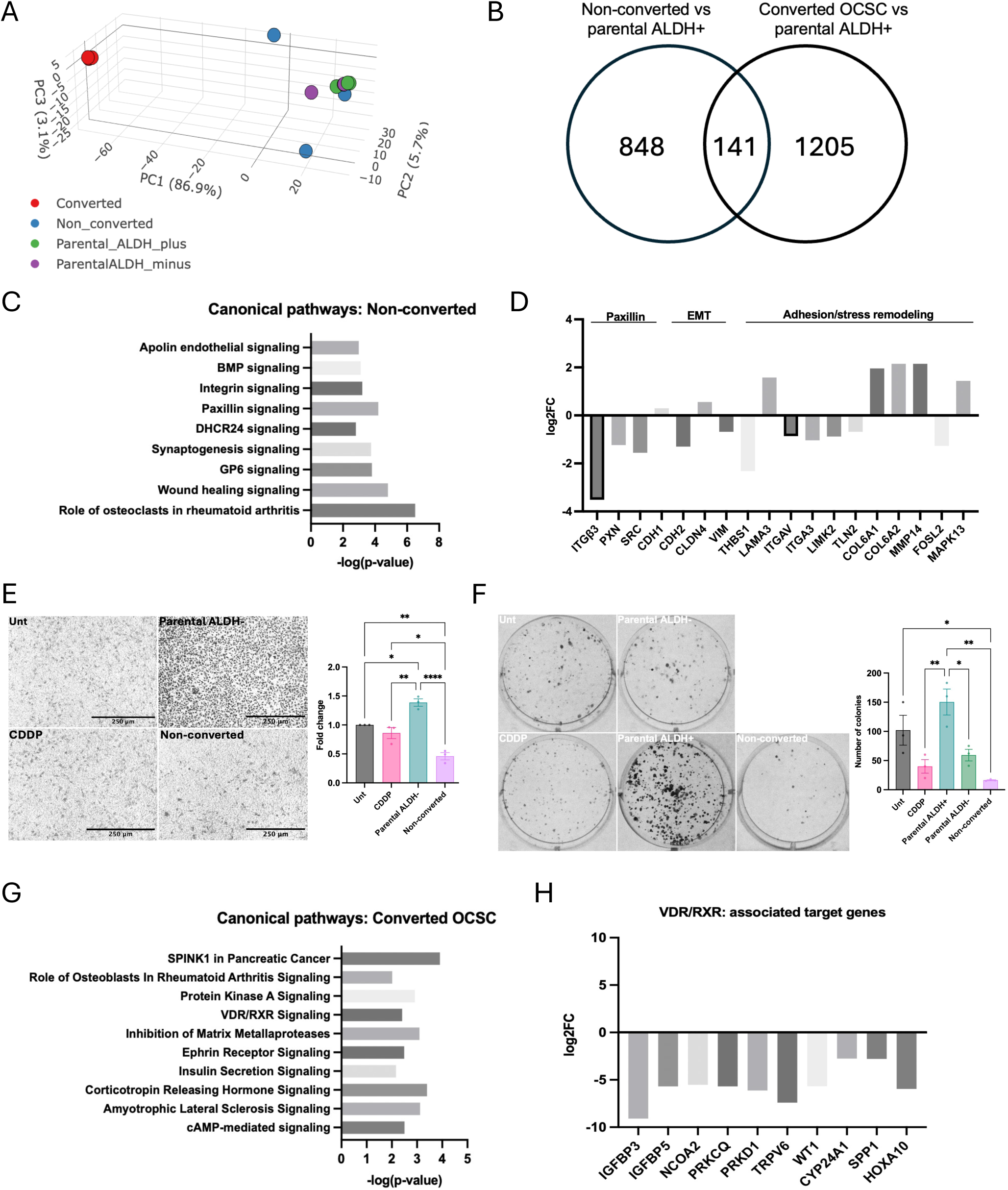
A. Principal component analysis (PCA) plot of parental ALDH+, parental ALDH-, non-converted and converted OCSCs. **B.** Venn diagram of overlapping genes between non-converted and converted OCSCs from RNA-seq analysis compared to parental ALDH+ cells. **C.** Ingenuity Pathway Analysis (IPA) of canonical pathways enriched in non-converted cells compared to parental ALDH+ cells. **D.** Fold change of gene expression from paxillin, EMT and adhesion/stress remodeling pathways in **(C)** using the IPA canonical pathways tool. **E.** Representative images of OVCAR5 untreated and cisplatin (16h, 15µM) treated whole cell populations and parental ALDH- and non-converted cells (5x10^5^) seeded in the top of the transwell insert and allowed to migrate for 24h and compared to untreated whole cell population. Quantification: right. Scale bar, 250µm. **F.** Representative images of OVCAR5 untreated and cisplatin (16h, 15µM) treated cells, parental ALDH+, parental ALDH- and non-converted cells (1.5x10^3^) seeded onto 6-well plates for colony formation. Cells were allowed to grow for 14 days before being imaged. Quantification: right. **G.** IPA analysis of canonical pathways enriched in converted OCSCs compared to parental ALDH+ cells. **H.** Fold change of gene expression from VDR/RXR pathway in **(G)** using the IPA canonical pathways tool. All experiments were performed in biological triplicate unless stated otherwise, +/- SEM, P *<0.5, P **<0.01, P ***<0.0001, P ****<0.00001.

Further analysis of non-converted cells revealed enrichment of apolin endothelial, BMP, integrin, paxillin and DHCR24 signaling pathways (Fig. 2C). Paxillin (PXN) signaling was explored further, as PXN is a central focal adhesion adaptor that integrates integrin/FAK/Src signaling to regulate cytoskeletal remodeling, cell migration, and EMT-associated programs across cancers^36^. Multiple genes involved in adhesion and cell remodeling were downregulated in non-converted cells, including PXN, SRC and ITGB3 as representative genes for focal adhesion signaling axis. Increased expression of known epithelial markers such as CDH1 and CLDN4 and decreased expression in mesenchymal markers CDH2 and VIM were also observed in non-converted cells (Fig. 2D). Functionally, non-converted cells had decreased migration and colony formation compared to cisplatin treated and parental ALDH-cells (Fig. 2E, F). To assess the clinical relevance of the 848 non-converted genes, RNA-seq data was compared to HGSC scRNA-seq datasets from treatment naïve (sensitive vs resistant) and NACT patients^37^ (Supplemental Fig. S4A). Cluster identification revealed 16 distinct clusters, with annotations for 14 of the clusters (Supplementary Fig. S4B, C). Within epithelial cells, 714 genes were identified between sensitive vs resistant patients and 972 genes in the naïve vs NACT patients (FC> |2|, FDR<0.05). Overlaying the 848 non-converted genes onto scRNA-seq data revealed the highest mean expression of upregulated genes in clusters 0 and 8 and downregulated genes in clusters 3, 5 and 10 (Supplementary Fig. S4D, E). Feature plots demonstrated the expression of SRC, PXN, ITGB3 and CADM1 (associated paxillin adhesion protein) across all clusters, with CADM1 expression highest in platinum-sensitive cluster 6 (Supplementary Fig. S4F). Overlapping genes between patients and non-converted cells in both datasets were identified (non-converted vs sensitive vs resistant: 23; non-converted vs naïve vs NACT: 27) (Supplementary Fig. S4G; Supplementary Table S2), including six common to all three comparisons, CLCNKA, ETV1, HCAR1, NEIL1, TRIM55 and ZNF385C (Supplementary Table S2). Overall, non-converted cells displayed an epithelial phenotype characterized by altered focal adhesion signaling and transcriptional similarities to HGSC patient cohorts.

Analysis of converted OCSCs identified SPINK1 pathway, protein kinase A signaling, VDR/RXR signaling and inhibition of matrix metalloproteases pathways (Fig. 2G). VDR/RXR (vitamin D receptor/retinoid X receptor) signaling was selected for further investigation, due to its role in cancer and its use of RXR as a shared binding partner with retinoic acid receptors (RAR) which have been linked to ALDH^38, 39^. Further analysis into VDR/RXR revealed downregulation of many VDR/RXR associated genes in converted OCSCs (Fig. 2H). Comparison of the 1205 converted OCSC genes with the same apteint datasets^37^ identified 737 and 811 differentially expressed epithelial genes between sensitive vs. resistant patients and 811 for naïve vs NACT patients, respectively (FC> |2|, FDR<0.05). Genes from converted OCSCs overlapped with both patient datasets (Supplementary Fig. S4H; Supplementary Table S3), with upregulated genes predominantly in NACT-associated clusters 9 and 16, and downregulated genes in platinum-sensitive clusters 2 and 6 (Supplementary Fig. S4I, J). Stemness scoring was calculated using GSEA curated datasets and identified clusters 3 (“stemness/stress response” ) and 10 (“secretory/stress response”), had the highest stemness scores, representing resistant and NACT patients respectively (Supplementary Fig. S4C, K). Feature plots showed prominent expression of POU5F1and PROM1 in resistant clusters (Supplementary Fig. S4L), while nuclear receptors RARA, RXRA and VDR were exhibited low to moderate expression across clusters (Supplementary Fig. S4L). Together, these findings demonstrate the clinical relevance of the converted OCSC transcriptional signature, which overlaps with epithelial populations from HGSC patients, particularly those associated with platinum resistance and NACT.

### Vitamin D decreases platinum-induced stemness

Given the downregulation of VDR/RXR associated genes in converted OCSCs, vitamin D was used to test whether activation of VDR/RXR signaling under vitamin D treatment could reduce stemness and inhibit platinum-induced conversion. Cells were treated with CDDP (IC_50_, 16h), vitamin D (IC_50_, 24h), or in combination (Supplementary Fig. S5A, B). Vitamin D alone decreased the ALDH+ population, while combination treatment significantly decreased both the ALDH+ population in bulk cells and the CDDP-induced conversion of sorted ALDH-cells (Fig. 3A). Vitamin D also reduced spheroid formation as well as the CDDP-induced increase in spheroid number in bulk and sorted cells (Fig. 3B; Supplementary Fig. S5C). Activation (increased expression of VDR) by vitamin D was confirmed by western blot (Fig. 3C, Supplementary Fig. S5D) and RT-qPCR for expression of VDR (Fig. 3D; Supplementary Fig. S5E). Additionally, vitamin D treatment decreased expression of stemness genes (BMI1, NANOG, OCT4, SOX2) compared to CDDP treatment alone, and combination treatement significantly decreased the CDDP-induced increase in stemness gene expression (Fig. 3D). Proximity ligation assay (PLA) was performed to determine pathway activation. Enhanced VDR:RXRα interactions following vitamin D treatment were observed by PLA (Fig. 3E; Supplementary Fig. S5F). Overall, vitamin D treatment activates VDR/RXR signaling and suppresses CDDP-induced stemness and conversion to ALDH+ cells.

**Figure 3.**
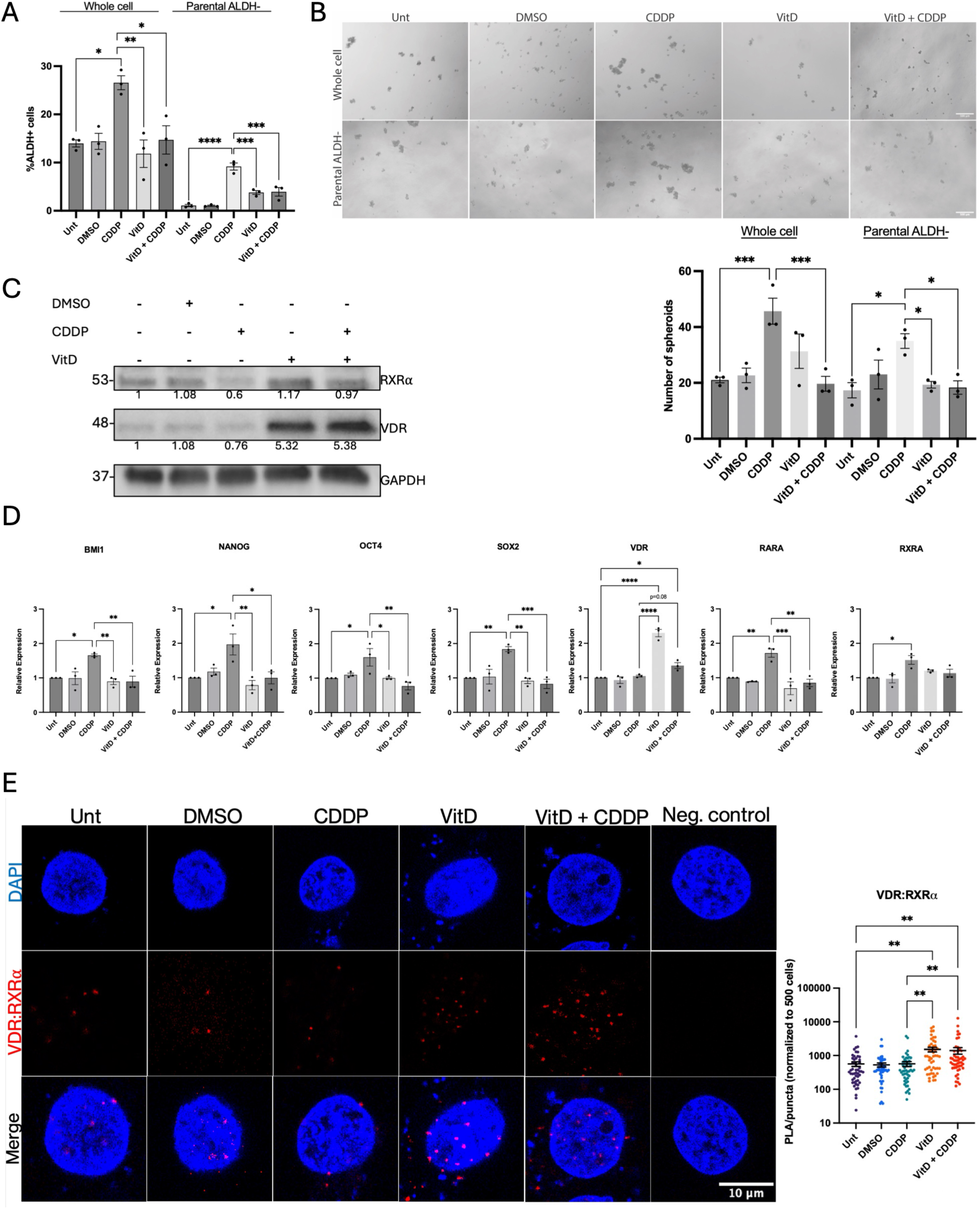
A. OVCAR3 cells (1x10^6^) treated with cisplatin (16h, 15µM), vitamin D (24h, 56nM) and combination treated and the ALDEFLUOR assay was performed to determine the percent of ALDH+ cells. Similar treatment scheme was performed on parental ALDH-cells. **B.** OVCAR3 cells treated with cisplatin (3h, 6µM), vitamin D (3h, 56nM) and combination treated and 2000 cells/well were seeded in low adhesion conditions after treatment. Parental ALDH-cells treated in a similar manner. Representative images of spheroid formation and quantification (>100um) (below). Scale bar, 500µm. **C.** OVCAR3 cells treated with cisplatin (16h, 15µM), vitamin D (24h, 56nM) and combination treated. Cells lysates were collected and analyzed for indicated proteins by western blot analysis. GAPDH was used as a loading control. **D.** Expression of stemness associated genes was determined by RT-qPCR analysis in cisplatin (16h, 15µM), vitamin D (24h, 56nM) and combination treated cells. **E.** Immunofluorescence representative images of VDR:RXR interactions in OVCAR3 cells by proximity ligation assay (PLA) after cisplatin (16h, 15µM), vitamin D (24h, 56nM) and combination treatments. Quantification: right. Scale bar, 10µm. All experiments were performed in biological triplicate unless stated otherwise, +/- SEM, P *<0.5, P **<0.01, P ***<0.0001, P ****<0.00001.

### Vitamin D decreases tumor growth in vivo

Based on our in vitro findings, it was of interest to test whether vitamin D could reduce growth in vivo. OVCAR3 cells (5x10^6^) were injected subcutaneously into female NSG mice, and once tumors reached 100mm^3^ were randomized into treatment groups (n=9, mice per group) of cisplatin (2mg/kg, IP) once weekly, vitamin D (0.5µg/kg, IP) three times weekly or combination of both treatments for three weeks (Fig. 4A). Body weight and tumor volume were monitored weekly throughout the duration of the study (Supplementary Fig. S6). As shown in Figure 4B, single drug treatments decreased tumor volume compared to control tumors; however, the combination of cisplatin and vitamin D resulted in the greatest reduction of tumor volume, with mean tumor volume staying relatively small throughout the duration of the study (<155mm^3^). At the end of the study, tumors from each treatment group were collected for IHC. IHC analysis of tumors showed vitamin D treatment significantly increased VDR expression (Fig. 4C), consistent with activation of the VDR signaling axis. In contrast, CDDP treatment significantly increased ALDH1A1, RARα and RXRα expression. Notably, the CDDP-induced increase in ALDH1A1, RARα and RXRα expression was attenuated when vitamin D was included in the combination treatment (Fig. 4C), consistent with our in vitro findings. In addition, CDDP increased expression of stemness-associated genes, whereas vitamin D treatment alone and in combination with CDDP was associated with reduced expression of these markers (Fig. 4D).

**Figure 4.**
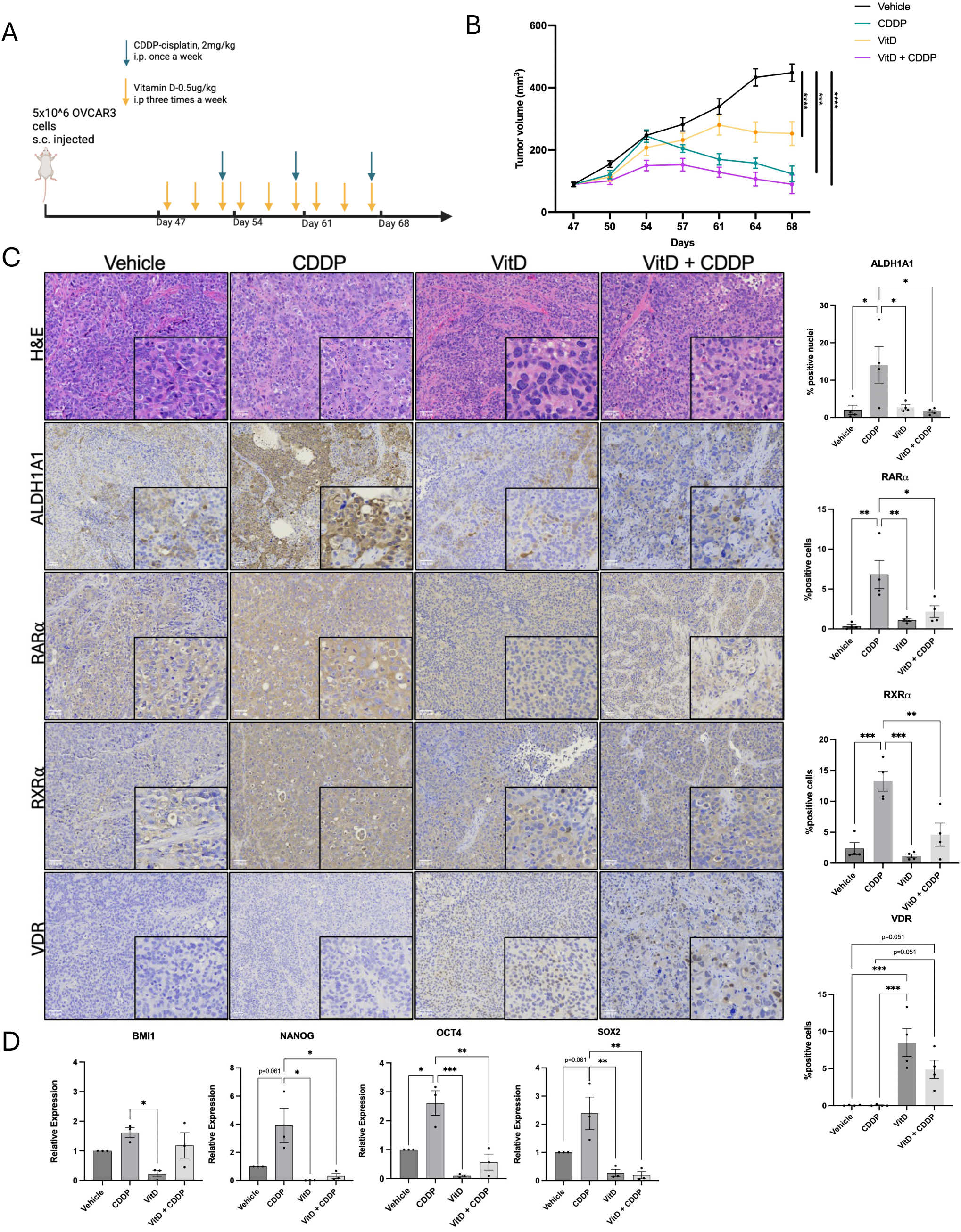
A. Treatment scheme for in vivo study. **B.** Mean tumor volume over the course of treatment in untreated, cisplatin, vitamin D and combination groups. **C.** Representative immunohistochemical (IHC) staining of ALDH1A1, RAR, RXR, VDR in tumors collected at the end of study, with corresponding quantification shown on the right (n=4), Scale bar, 100µm. **D.** Expression of stemness associated genes was determined by RT-qPCR analysis in untreated, cisplatin, vitamin D and combination treated tumors. All experiments were performed in biological triplicate unless stated otherwise, +/- SEM, P *<0.5, P **<0.01, P ***<0.0001, P ****<0.00001.

### Platinum increases RAR/RXR activity

Because ALDH activity contributes to atRA synthesis, we hypothesized that CDDP-induced increase in ALDH activity would result in elevated atRA levels. To test this, ELISA was performed on cells treated with CDDP for 16h. After CDDP treatment, the atRA level significantly increased (Fig. 5A). Consistent with pathway activation, CDDP treatment increased RARα protein levels and RARα and RXRα at mRNA levels, along with expression of canonical RAR/RXR target genes, CYP26B1 and STRA6^40, 41^ (Fig. 5B, C; Supplementary Fig. S7A). In addition, PLA demonstrated an increase in RARα:RXRα interactions following CDDP and atRA treatments compared to untreated cells (Fig.5D). Increased RARα:RXRα interactions were also observed in parental ALDH+ and converted OCSCs compared to parental ALDH- and non-converted cells (Fig. 5D; Supplementary Fig. S7B). To further confirm that CDDP activated RAR/RXR signaling, cells transfected with RARE-luciferase reporter were treated with CDDP or atRA. Both treatments significantly increased RAR/RXR activity compared to untreated cells and cells treated with RAR/RXR activity inhibitor, AGN194310 (10µM, 24h) (Fig. 5E).

**Figure 5.**
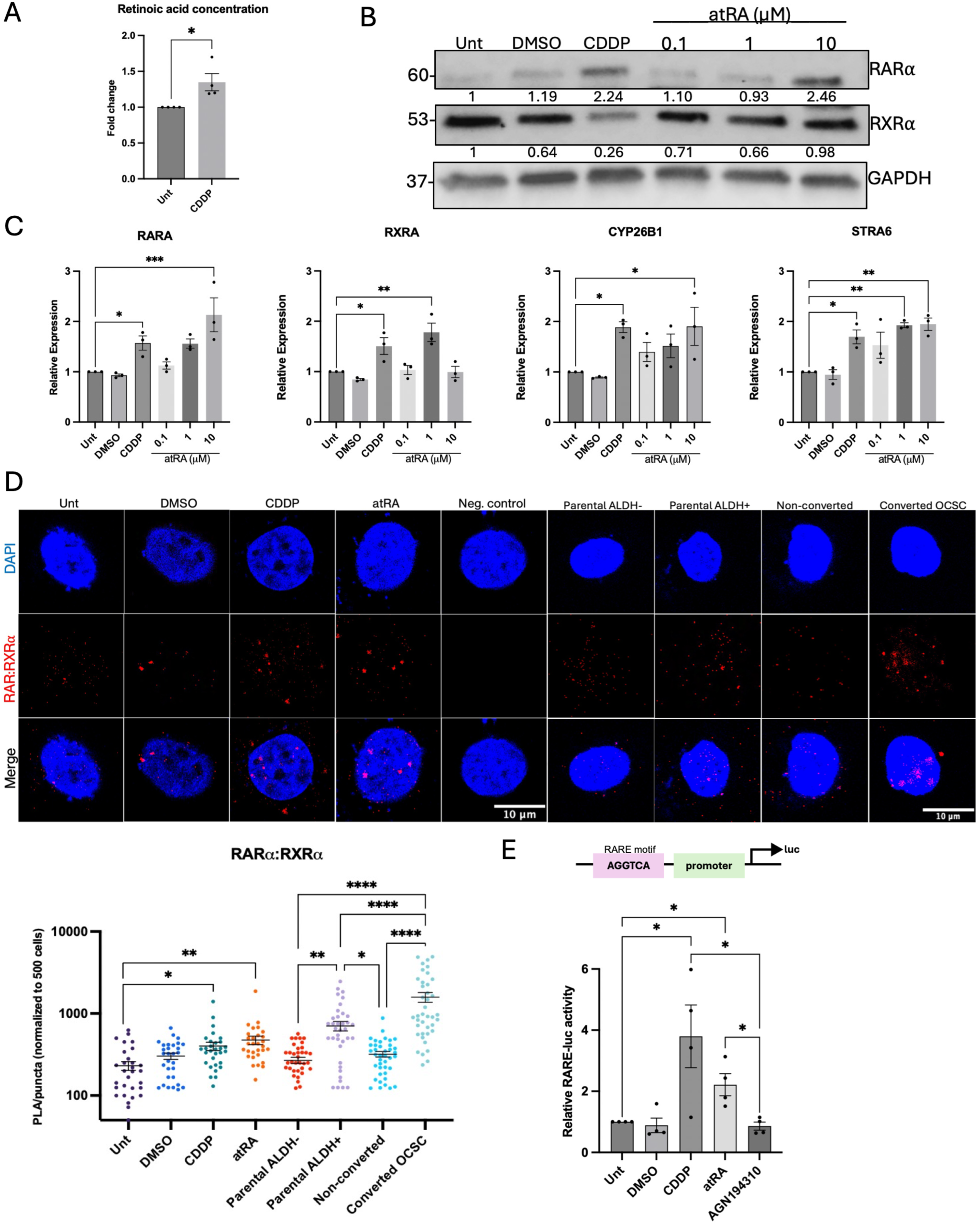
A. Intracellular levels of all-trans retinoic acid (atRA) were measured by ELISA in OVCAR3 untreated and cisplatin (16h, 15µM) treated cells. **B.** OVCAR3 cells treated with indicated doses of atRA for 24h. Cell lysates were collected and analyzed for indicated proteins by western blot analysis. GAPDH was used as a loading control. **C.** Expression of atRA signaling receptors and downstream target genes after indicated doses of atRA (24h). **D.** Immunofluorescence representative images of RAR:RXR interactions in OVCAR3 cells after cisplatin (16h, 15µM) or atRA (24h, 10µM) treatments and sorted cell populations by PLA. Quantification: below. **E.** OVCAR5 cells were transfected with RARE-luc reporter and a Renilla luciferase plasmid (24h). Cells were treated with cisplatin (16h, 15µM), atRA (24h, 10µM) or RAR/RXR antagonist, AGN194310 (24h, 10µM). Luciferase activity was measured, normalized to Renilla luciferase and compared to untreated cells. All experiments were performed in biological triplicate unless stated otherwise, +/- SEM, P *<0.5, P **<0.01, P ***<0.0001, P ****<0.00001.

We next investigated the relationship between ALDH activity and CDDP-induced RAR/RXR signaling. Cells were treated with increasing doses of ALDH activity inhibitor, DEAB (25-100μM, 72h) and analyzed by western blot. DEAB treatment reduced expression of RARα, RXRα, and the RAR/RXR signaling co-activator, NCOA2^42^ (Supplementary Fig. S8A). To confirm that DEAB effectively suppressed atRA levels, ELISA was performed on cells after DEAB treatment (75µM, 72h), showing a reduction in atRA. (Supplementary Fig. S8B). To assess the impact of atRA depletion on receptor complex formation, PLA was performed on cells treated with DEAB in combination with CDDP or atRA. CDDP and atRA increased RARα:RXRα interactions relative to DEAB treatment alone (Supplementary Fig. S8C, columns 3-5); however, co-treatment with DEAB suppressed interactions even in the presence of activating ligands (Supplementary Fig. S8C, columns 6, 8). In contrast, removal of DEAB prior to CDDP or atRA treatments restored RARα:RXRα interactions (Supplementary Fig. S8C, columns 7, 9). Together, these findings indicate that inhibition of ALDH activity reduces atRA levels and attenuates CDDP and atRA associated RARα:RXRα complexes.

### Vitamin D suppresses RAR/RXR activity and alters RXR partner engagement

Based on previous results that vitamin D decreased stemness, we next tested whether vitamin D could alter RXR partner engagement in the presence of CDDP and shift towards VDR/RXR partner engagement. Cells were treated with vitamin D alone or in combination with atRA and CDDP, and expression of RARA, RXRA, VDR and downstream RAR/RXR genes (CYP26B1, STRA6) was measured. Similar to previous results, CDDP and atRA increased RARA and RXRA whereas vitamin D treatment increased VDR expression (Fig. 6A). Addition of vitamin D reduced expression of RAR/RXR target genes, with decreased CYP26B1 expression following atRA + vitamin D treatment and decreased STRA6 expression following both atRA + vitamin D and CDDP + vitamin D treatments (Fig 6A). Expression of each receptor, measured by western blot also revealed a similar pattern where combination treatments with vitamin D, decreased RARα expression relative to CDDP and atRA single treatments (Fig 6B). Furthermore, RAR/RXR transcriptional activity was measured using the same RARE-luciferase reporter assay. As expected, treatment with CDDP and atRA increased RAR/RXR activity and vitamin D treatment had minimal activity (Fig. 6C). Combination treatments with vitamin D also showed a reduction in RAR/RXR activity even when CDDP and atRA were present (Fig. 6C). Similarly, PLA demonstrated that vitamin D treatment increased VDR:RXRα interactions while reducing RARα:RXRα interactions whenever vitamin D was present (Fig. 6D, Supplemental Fig. S9). These findings show that vitamin D treatment is associated with reduced RAR/RXR signaling and increased VDR:RXRα complex formation, consistent with altered RXR partner engagement.

**Figure 6.**
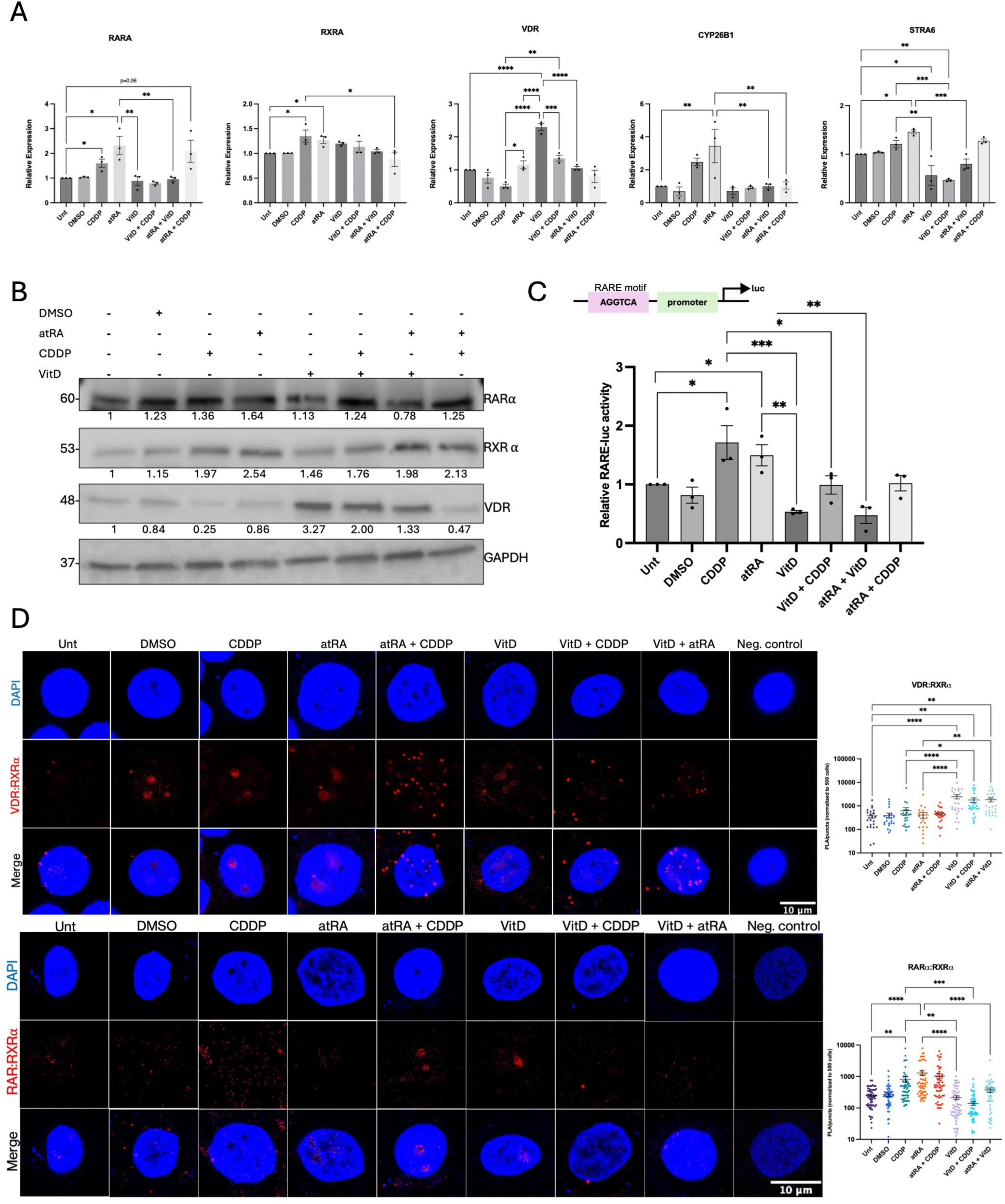
A. Expression of nuclear signaling receptors and downstream RAR/RXR target genes after cisplatin (16h, 15µM), atRA (24h, 10µM), or vitamin D (24h, 56nM) treatments. **B.** OVCAR3 cells treated with cisplatin (16h, 15µM), atRA (24h, 10µM), or vitamin D (24h, 56nM). Cell lysates were collected and analyzed for indicated proteins by western blot. **C.** OVCAR5 cells were transfected with RARE-luc reporter and a Renilla luciferase plasmid. Cells were treated with CDDP (16h, 15µM), atRA (24h, 10µM) or vitamin D (24h, 56nM). Luciferase activity was measured, normalized to Renilla luciferase and compared to untreated cells. **D.** Top: Immunofluorescence representative images of VDR:RXRα interactions in OVCAR3 cells by PLA after cisplatin (16h, 15µM), atRA (24h, 10µM), or vitamin D (24h, 56nM) treatments. Below: Immunofluorescence representative images of RARα:RXRα interactions using the same treatment scheme. Quantification: right. Scale bar, 10µm. All experiments were performed in biological triplicate unless stated otherwise, +/- SEM, P *<0.5, P **<0.01, P ***<0.0001, P ****<0.00001.

## Discussion

Platinum resistance and tumor recurrence remain major challenges in treatment of HGSC. Recent evidence demonstrates that CSC plasticity provides a mechanism for survival and subsequent expansion of this subpopulation of cells following chemotherapy^14, 43^. However, a role for cell plasticity in stress-induced conversion of a non-CSC to CSC remains to be fully established. Here, we provide evidence that platinum treatment promotes OCSC plasticity by inducing the conversion of non-OCSCs into OCSCs with increased stemness properties. Transcriptomic analysis of converted OCSCs identified VDR/RXR as a potential pathway associated with plasticity, prompting investigation into VDR and its heterodimeric partner, RXR. Examination of RXR-associated nuclear receptor RAR revealed that platinum treatment increased RAR/RXR activity, whereas activation of VDR/RXR with vitamin D inhibited stemness, blocked platinum-induced conversion of non-OCSCs to OCSCs and shifted RXR partner utilization toward VDR. These findings suggest that the differential engagement of RXR to binding partners RAR and VDR may influence OCSC states following platinum exposure. Consistent with these findings, vitamin D in combination with platinum decreased tumor formation in vivo. Collectively, our study identifies vitamin D as a potential therapeutic strategy to limit therapy-induced OCSC plasticity to prevent recurrence.

Our initial findings demonstrated that platinum treatment not only increased the OCSC population but also promoted the conversion of ALDH-cells to ALDH+ cells. Importantly, sorted ALDH-cells remained phenotypically stable in the absence of treatment, supporting a treatment-driven transition and not a stochastic conversion. Converted OCSCs exhibited enhanced stem-like properties, indicating that these cells acquired functional stem-like properties and that platinum-induced conversion generates a distinct stem-like state compared to pre-existing CSCs. Previous studies have demonstrated CSC plasticity can be induced in response to chemotherapeutic agents, where cells not only retain their stemness features, but also exhibit tumor formation capabilities in vivo compared to non-CSCs^44–46^. Our findings are consistent with these reports stating that CSC populations are not solely maintained through expansion of pre-existing CSCs or depletion of non-CSCs but can also arise through transitions between different cellular states^45, 47^.

Transcriptomic analysis revealed distinct molecular programs between non-converted cells and converted OCSCs following platinum treatment. Non-converted cells were enriched for pathways associated with integrin signaling, EMT and adhesion/stress remodeling, processes previously implicated in CSC plasticity and transitional cellular states^48, 49^. EMT is a key mechanism underlying CSC plasticity, as transitions between epithelial and mesenchymal states contribute to shifts between non-CSC and CSC phenotypes. Adhesion remodeling can accompany these transitions and reflects cytoskeletal and extracellular matrix adaptations that occur during these transitions^18, 50–54^. Reduced expression of PXN signaling components together with increased expression of epithelial markers suggests that non-converted cells retained a more epithelial and differentiated phenotype, despite platinum exposure. These findings also suggest that while platinum induced cellular stress and remodeling programs, non-converted cells remained largely in a non-CSC state, representing an intermediate, chemotherapy induced phenotype, consistent with previously described partial or hybrid EMT states associated with CSC plasticity^55–57^. The presence of similar transcriptional features in therapy-associated tumor states from scRNA-seq^37^ datasets further support the clinical relevance of this intermediate population and suggests that these cells could represent markers for cells poised for CSC conversion. In contrast, repression of VDR/RXR signaling in converted OCSCs suggested loss of differentiation-associated transcriptional programs during acquisition of stemness. Previous studies have demonstrated that vitamin D treatment suppresses stemness through inhibition of stemness-related pathways including, Wnt, NF-κB and NOTCH and by binding to vitamin D response elements (VDRE) on stemness genes such as SOX2^58–64^. Clinical studies have also associated vitamin D supplementation with reduced cancer mortality^65, 66^. Given the established role of vitamin D in promoting differentiation, we found that vitamin D reduced stemness properties of cells, decreased the OCSC population and suppressed the platinum-induced conversion of non-OCSCs into OCSCs. Together, these findings suggest that vitamin D signaling can function to suppress therapy-induced stemness and cellular plasticity. Furthermore, comparison of the converted OCSC signature with scRNA-seq datasets^37^ revealed overlap with resistant and NACT-treated patient populations, supporting the translational relevance of the converted phenotype in patients. These effects were also observed in vivo where vitamin D and CDDP treatments decreased tumor volume alone, but combination treatment had the greatest reduction in tumor growth, supporting the antitumor efficacy of vitamin D. IHC analysis demonstrated that vitamin D attenuated CDDP-induced increase in RARα and RXRα expression, demonstrating suppression of RAR/RXR signaling components. The opposing changes in RARα and VDR, despite no significant change in RXRα for western analysis, suggest that treatment response may depend more on RXR heterodimerization and pathway activity than on RXR abundance alone^67–69^. Furthermore, decreased expression of stemness-associated genes after vitamin D treatment demonstrated that even after continuous vitamin D treatment expression was attenuated.

Because ALDH enzymes catalyze oxidation of retinaldehyde to atRA^70^ our findings demonstrate that platinum treatment activates an ALDH-dependent increase in atRA, resulting in increased RAR/RXR signaling. The dependence of RAR/RXR activation of ALDH-mediated atRA production supports a mechanistic link between increased ALDH activity and activation of the pathway following platinum treatment. Elevated RARα:RXRα interactions after platinum treatment and in parental ALDH+ and converted OCSCs identifies RAR/RXR signaling as a feature associated with stemness. Previous studies have demonstrated that RAR/RXR signaling exerts context-dependent effects, promoting either differentiation or stemness depending on cellular state and signaling environment, highlighting its involvement in plasticity^28, 71–73^.

Perhaps one of the most significant findings of this study is that vitamin D altered RXR partner engagement even in the presence of activating RAR ligands. RXR serves as an obligate heterodimer partner for both VDR and RAR, allowing ligand availability, receptor abundance and cellular context to influence heterodimer formation and downstream transcriptional responses^24, 67, 68, 74, 75^. Consistent with this, vitamin D treatment attenuated RAR/RXR signaling while increasing VDR:RXRα interactions suggesting that activation of VDR redirects RXR partner engagement even during CDDP and atRA treatments. Importantly, because vitamin D treatment also reduced stemness and inhibited platinum-induced CSC conversion, these data suggest that modulation and balance of nuclear receptor signaling dynamics may contribute to blocking therapy-induced cellular plasticity.

Although these findings support a role for platinum in nuclear receptor signaling in OCSC plasticity, several limitations should be considered. First, conversion assays were performed after one platinum treatment and therefore may not show the effects multiple cycles of chemotherapy have on plasticity processes. Second, although platinum treatment increased RAR/RXR activity and converted OCSCs exhibited increased RARα:RXRα interactions, the present study did not determine whether RAR/RXR signaling is required for OCSC conversion. Rather it identifies RAR/RXR signaling as a pathway that is enhanced alongside the platinum-induced acquisition of the OCSC phenotype. Similarly, although vitamin D increased VDR:RXRα interactions while suppressing conversion, these experiments did not establish that altered RXR partner engagement was solely responsible for the observed phenotypic effects. Additional genetic or selective pharmacologic approaches targeting RAR^76, 77^, VDR^78^ or RXR isoforms will be needed to define the requirement of each pathway component during OCSC conversion.

In summary, our findings demonstrate that platinum treatment promotes OCSC plasticity through conversion of non-OCSCs to OCSCs, while vitamin D suppresses this process. Platinum treatment increased RAR/RXR signaling, while vitamin D increased VDR/RXR engagement, reduced RAR/RXR signaling and inhibited OCSC conversion and stemness. Together, these findings identify a balance between RAR/RXR and VDR/RXR signaling is associated with OCSC states following platinum exposure and identify nuclear receptor signaling as potential regulator of OCSC plasticity.

## Acknowledgements

We thank the Flow Cytometry Core Facility (Indiana University, Bloomington, IN) and Christiane Hassel for technical assistance with flow cytometry. We thank Dr. Heather M. O’Hagan (Indiana University School of Medicine) and Dr. Peter C. Hollenhorst for helpful discussions. We would like to thank the Light Microscopy Imaging Center (Indiana University, Bloomington, IN) and Andras Kun for assistance with confocal microscopy. We thank the shared facilities of the Indiana University Melvin and Bren Simon Comprehensive Cancer Center, including the Laboratory Animal Resource Center along with Kathy Coy, Melinda Ervin, Melissa Trowbridge, Sally Bosque, Tony Sinn, and especially Felicia M. Kennedy for her assistance with the in vivo studies. We thank the Center for Genomics and Bioinformatics (Indiana University, Bloomington, IN) for their assistance with RNA-seq, especially Ram Podicheti. With gratitude to the Jerry and Peggy Throgmartin Family for their philanthropic investments in our work.

