## Supplementary Tables for "Activation of vitamin D signaling suppresses platinum-induced ovarian cancer stem cell plasticity"

Supplementary Table S1. Primer sequences used for qRT-PCR

| **Primer** | **Forward (5’🡪3’)** | **Reverse (5’🡪3’)** |
| --- | --- | --- |
| EEF1A1 | TTGTCGTCATTGGACACGTAG | GATACCACGTTCACGCTCAG |
| ALDH1A3 | CTTCTGCCTTAGAGTCTGGAAC | CGTATTCACCTAGTTCTCTGCC |
| BMI1 | TTACCTGGAGACCAGCAAGT | CATTAGAGCCATTGGCAGCA |
| NANOG | CAGAAGGCCTCAGCACCTACCTACCC | AAACCGACCTTGACGTACGTCCTGAC |
| OCT4 | GGAAAGGCTTCCCCCTCAGGGAAAGG | AAGAACATGTGTAAGCTGCGGCCC |
| SOX2 | GCGCGGGCGTGAACCAG | CGGCGCGGGGAGATACA |
| RARA | CCAGCACCAGCTTCCAGTTA | GGCTTGTAGATGCGGGGTAG |
| RXRA | GACCTACGTGGAGGCAAACA | AGAAGTGTGGGATCCGCTTG |
| VDR | CCCTTCTGTGACCCTAGAGC | GCAGTACGATCTGGTCCTCA |
| CYP26B1 | CATCGGAGAGACCGGCCA | GCCAATGGAATTGGACACCG |
| STRA6 | CCATCTGTGGGCTCTGGAAG | TCGAAGGTTGGTCCTGTGTG |

Supplementary Table S2. Shared genes between non-converted cells and scRNA-seq datasets

| **Comparison** | **Number of genes in common** | **Gene** |
| --- | --- | --- |
| Non-converted vs sensitive vs resistant | 23 | APBA1, ATAD3C, CAPN12, CAPS, CHST2, ENOX1, ETV1, FGF9, FREM2, GPCR5AHCAR1, LGSN, LINC02609, LONRF2, NEIL1, RERG, ST6GALNAC5, TMEM74B, TP63, TRIM55, ZNF385C, ZNF518B |
| Non-converted vs naïve vs NACT | 27 | ADAMTS16, AFF2, AL162388.2, AMIGO2, AQP3, ARHGAP31, BDNF, CASKIN1, CEBPA, CLCKNA, COL8A1, CSF1R, ETV1, HCAR1, IL6, KRT15, LINC01235, LYNX1, MB, NEIL1, P3H2, PRTFDC1, SP6, SVOPL, TRIM55, TTC34, ZNF385C |
| Non-converted vs sensitive vs resistant and non-converted vs naïve vs NACT | 6 | CLCNKA, ETV1, HCAR1, NEIL1, TRIM55 and ZNF385C |

Supplementary Table S3. Shared genes between converted OCSCs and scRNA-seq datasets

| **Comparison** | **Number of genes in common** | **Gene** |
| --- | --- | --- |
| Converted OCSC vs sensitive vs resistant | 50 | ADAMTS1, ANOS1, APCDD2L-DT, AQP9, BRINP1, CA9, CCDC144, LA-AS1, CD274, CXCL1, DCC, DCLK1, DKK2, ERG, ERP27, FOXG1, GBP5,GJB2, HGD, KCNK2, LAD1, LERFS, LINC01606, LMO1, MAP7D2, MASP2, MMP12, MMP3, NPR3, PIGR, POU5F1, PRKG1, PROM2, PSG9, RAB25, S100A9, SAMD5, SCEL, SMIM22, SPINT1, SPOCK2, SPSB4, TLL1, TRHDE-AS1 |
| Converted OCSC vs naïve vs NACT | 51 | ADAMTSL13, ALX1, APRL14, C13orf46, CDH18, CHT1, CILP, CLEC7A, CRABP1, CRYAB, CSMD1, DDIT4L, DGKG, DMRT2, DOCK10, DSC3, ELAVL2, EPHA7, GALNT13, HCK, HFM1, HOXA9, IRX3, KRT17, KRT19, KRT4, LGI2, LINC00839, LINC001127, MILR1, NPR3, NRAD1, NTRK2, NYAP5, PRDM5, PRSS12, RGCC, RIMS1, RIMS4, SH3GL3, SOX2, TENM1, TMSB15A, TRPV2, UCHL1, VEGFC, ZBDBF2, ZFHX4 |
| Converted OCSC vs sensitive vs resistant and converted OCSC vs naïve vs NACT | 1 | NPR3 |
