## Supplementary Figures for "Activation of vitamin D signaling suppresses platinum-induced ovarian cancer stem cell plasticity"

### Slide 1
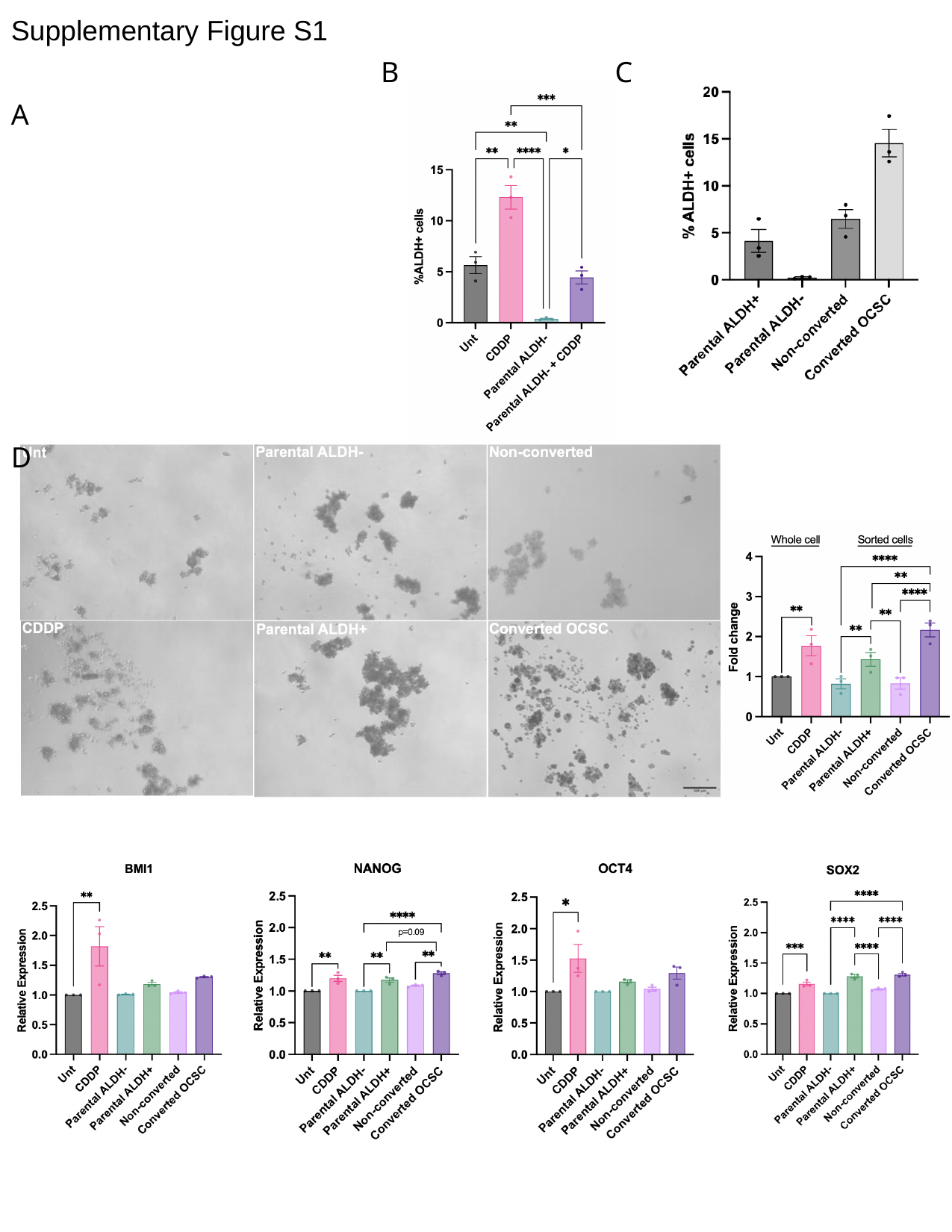

#### Supplementary Figure S1
B
C
A
D
E

### Slide 2
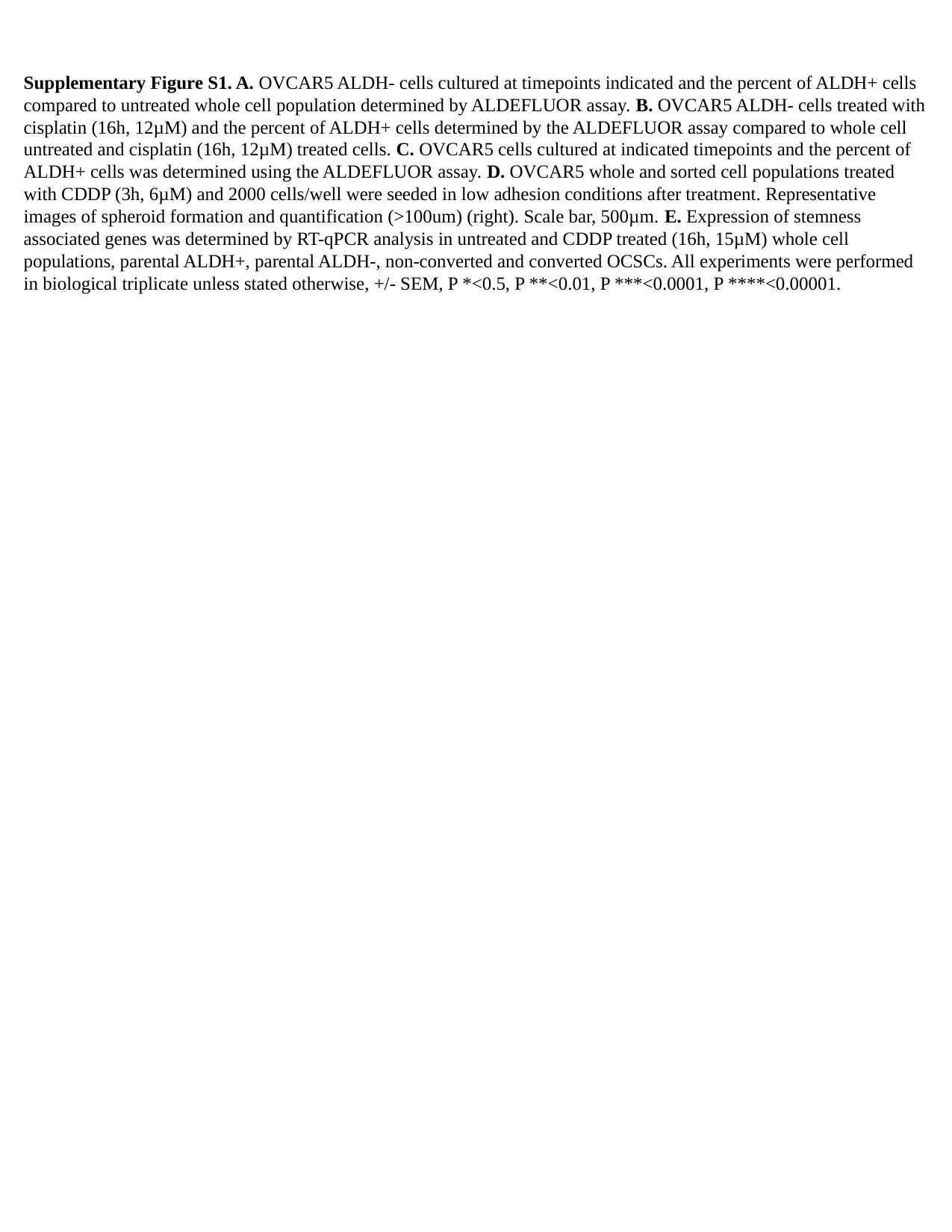

Supplementary Figure S1. A. OVCAR5 ALDH- cells cultured at timepoints indicated and the percent of ALDH+ cells compared to untreated whole cell population determined by ALDEFLUOR assay. B. OVCAR5 ALDH- cells treated with cisplatin (16h, 12µM) and the percent of ALDH+ cells determined by the ALDEFLUOR assay compared to whole cell untreated and cisplatin (16h, 12µM) treated cells. C. OVCAR5 cells cultured at indicated timepoints and the percent of ALDH+ cells was determined using the ALDEFLUOR assay. D. OVCAR5 whole and sorted cell populations treated with CDDP (3h, 6µM) and 2000 cells/well were seeded in low adhesion conditions after treatment. Representative images of spheroid formation and quantification (>100um) (right). Scale bar, 500µm. E. Expression of stemness associated genes was determined by RT-qPCR analysis in untreated and CDDP treated (16h, 15µM) whole cell populations, parental ALDH+, parental ALDH-, non-converted and converted OCSCs. All experiments were performed in biological triplicate unless stated otherwise, +/- SEM, P *<0.5, P **<0.01, P ***<0.0001, P ****<0.00001.

### Slide 3
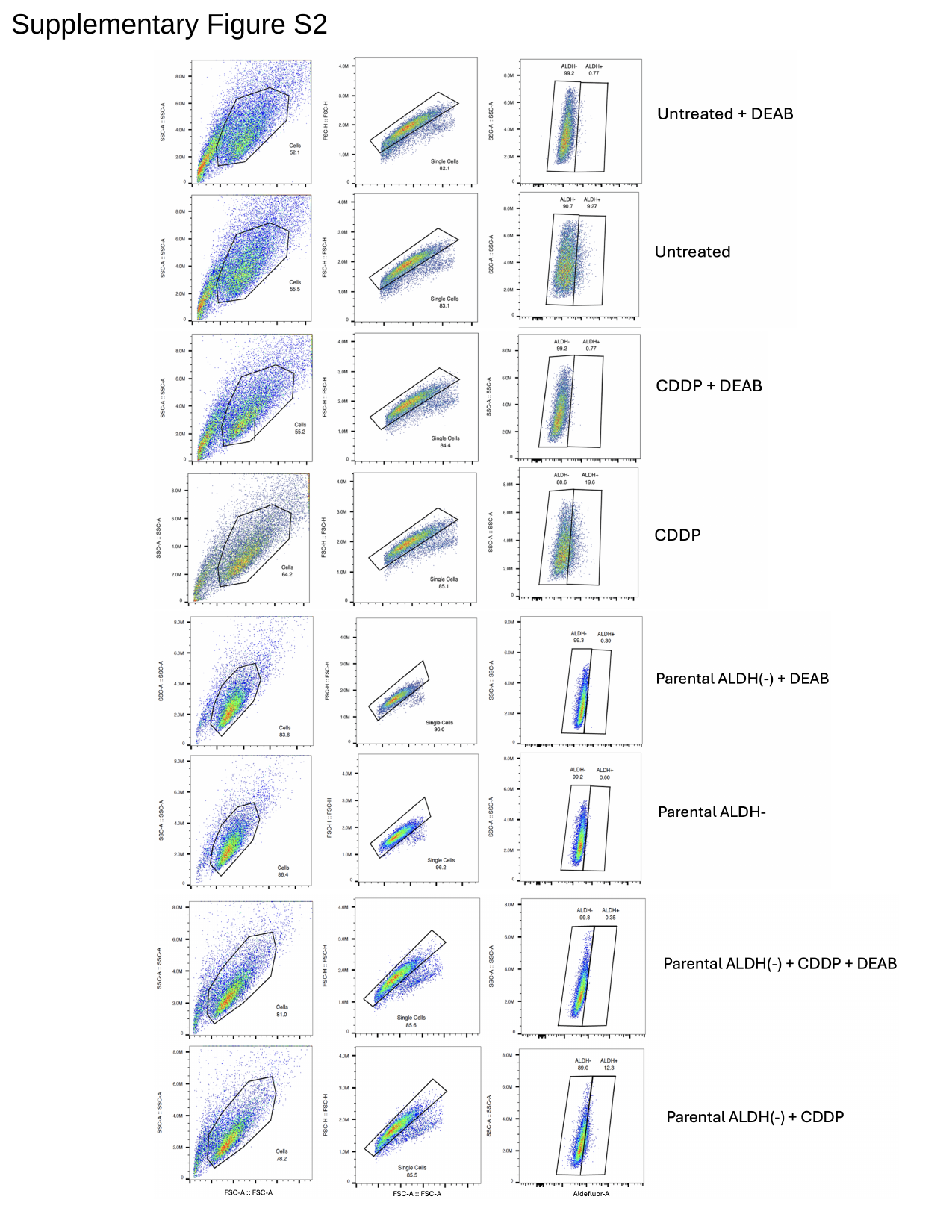

#### Supplementary Figure S2

### Slide 4
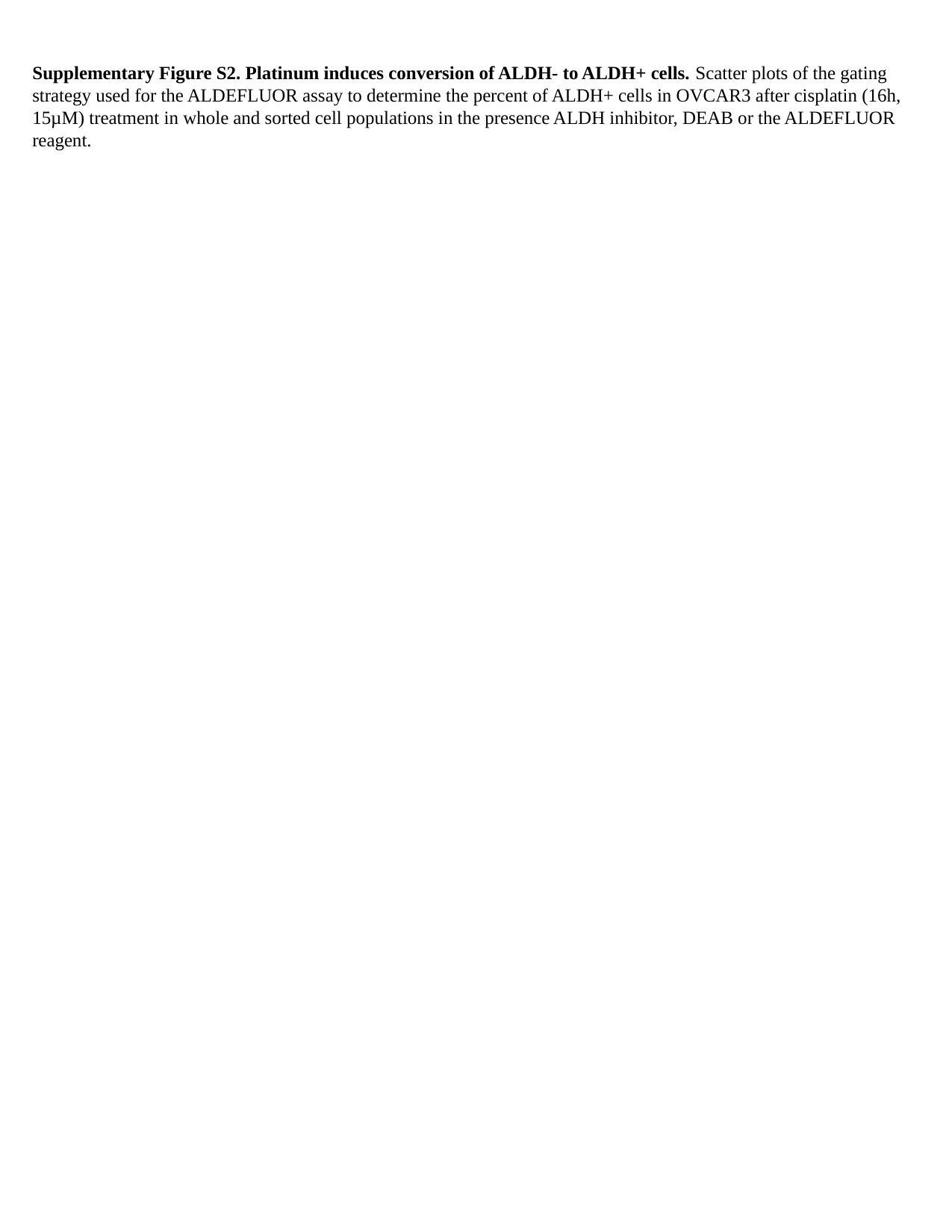

Supplementary Figure S2. Platinum induces conversion of ALDH- to ALDH+ cells. Scatter plots of the gating strategy used for the ALDEFLUOR assay to determine the percent of ALDH+ cells in OVCAR3 after cisplatin (16h, 15µM) treatment in whole and sorted cell populations in the presence ALDH inhibitor, DEAB or the ALDEFLUOR reagent.

### Slide 5
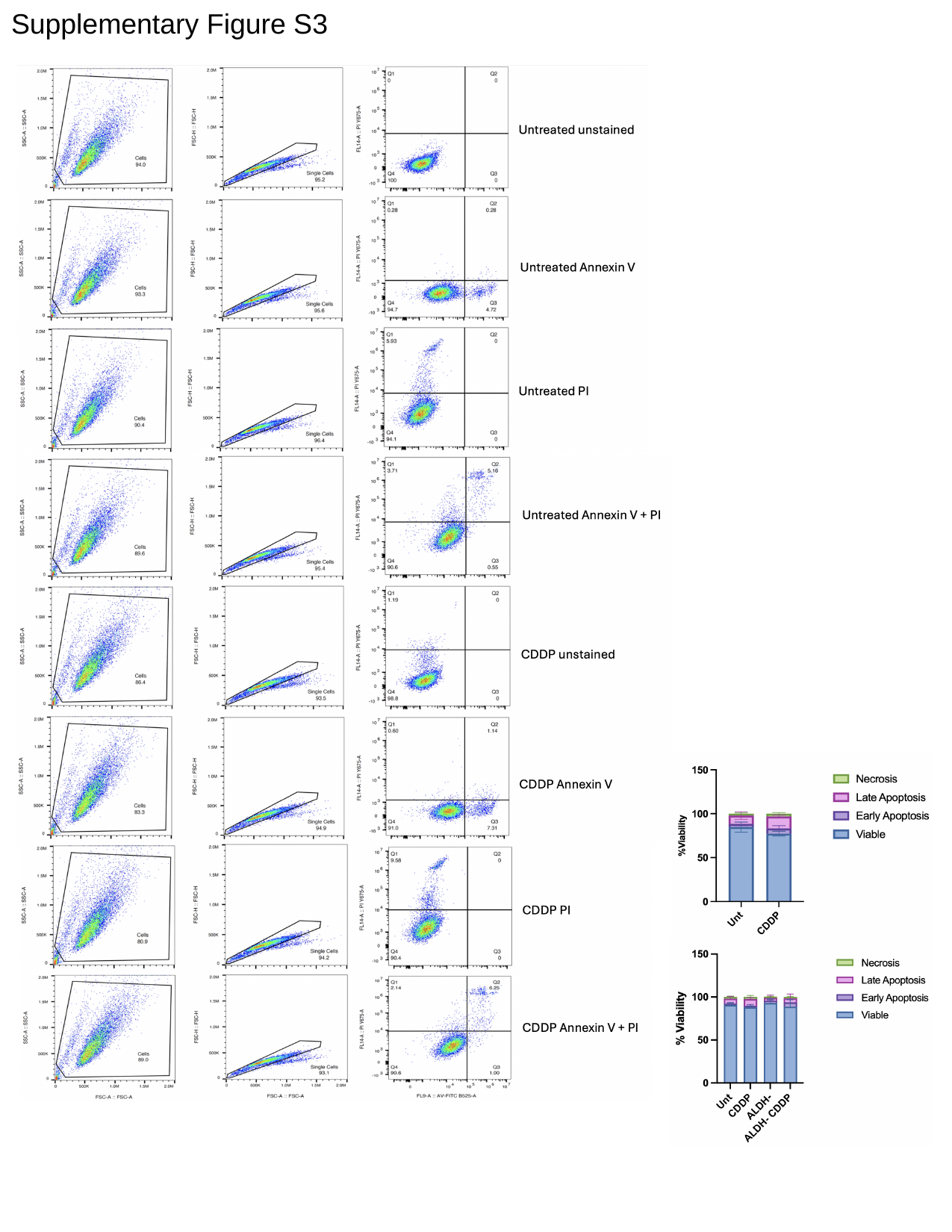

#### Supplementary Figure S3

### Slide 6
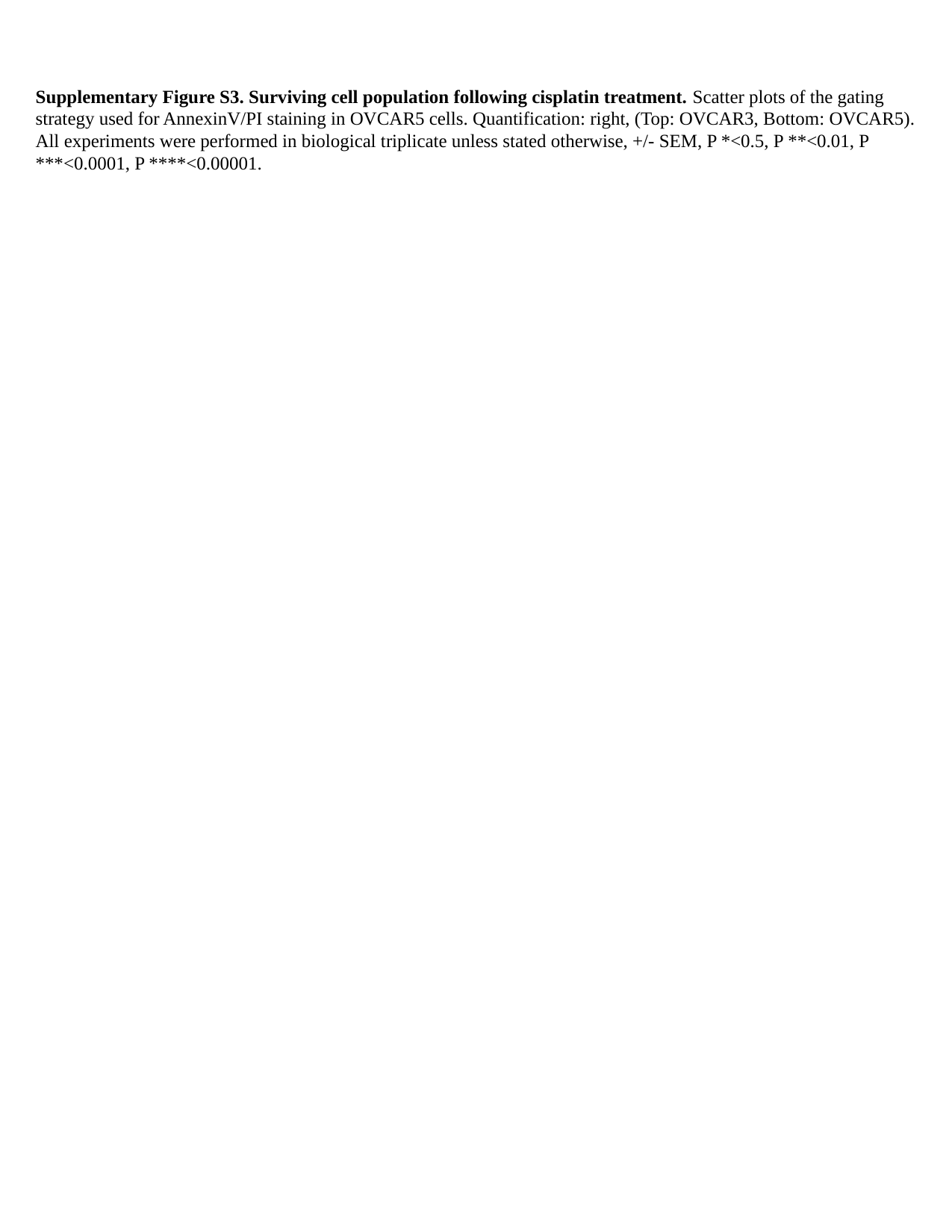

Supplementary Figure S3. Surviving cell population following cisplatin treatment. Scatter plots of the gating strategy used for AnnexinV/PI staining in OVCAR5 cells. Quantification: right, (Top: OVCAR3, Bottom: OVCAR5). All experiments were performed in biological triplicate unless stated otherwise, +/- SEM, P *<0.5, P **<0.01, P ***<0.0001, P ****<0.00001.

### Slide 7
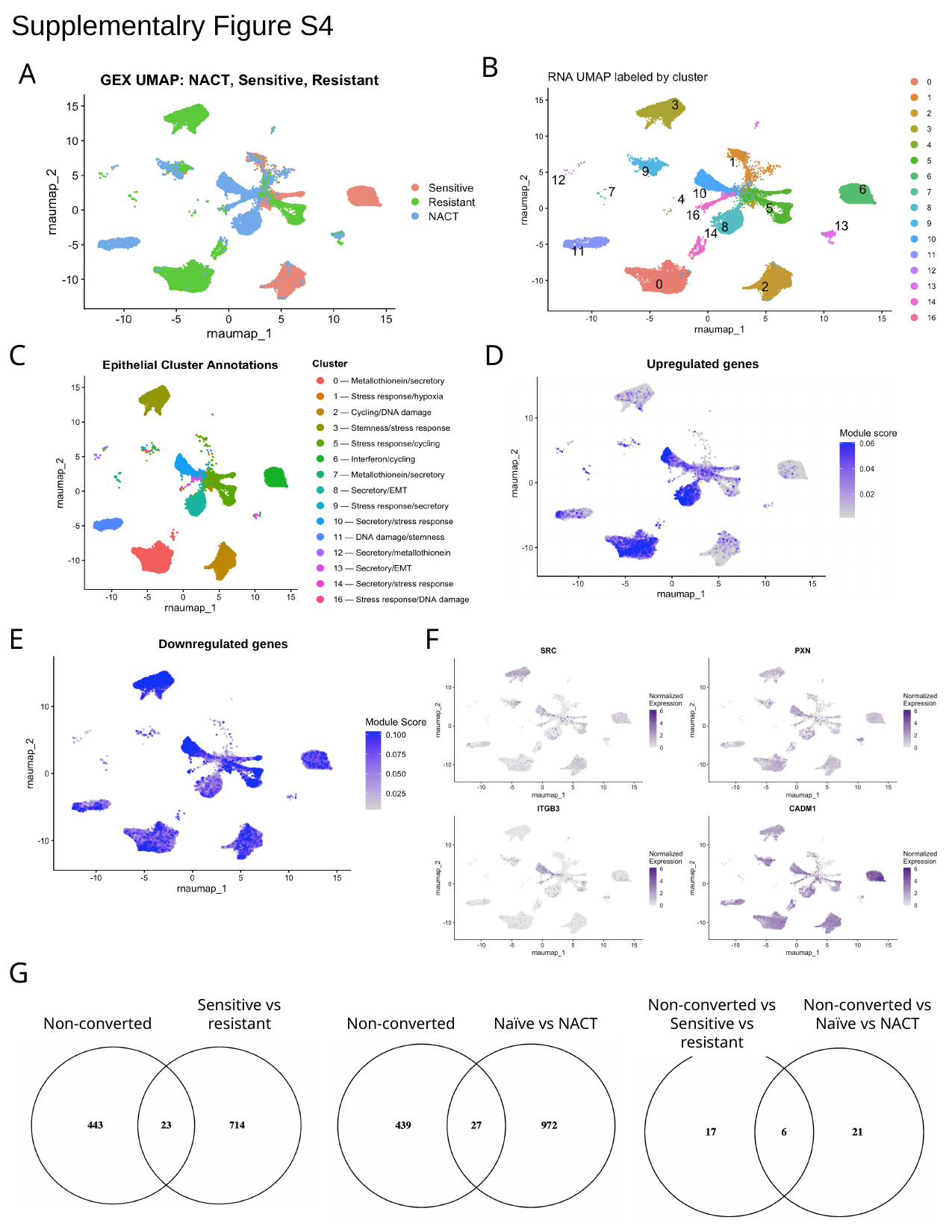

#### Supplementalry Figure S4
B
A
C
D
Upregulated genes
E
F
Downregulated genes
G
Sensitive vs resistant
Non-converted vs Sensitive vs resistant
Non-converted vs Naïve vs NACT
Non-converted
Non-converted
Naïve vs NACT

### Slide 8
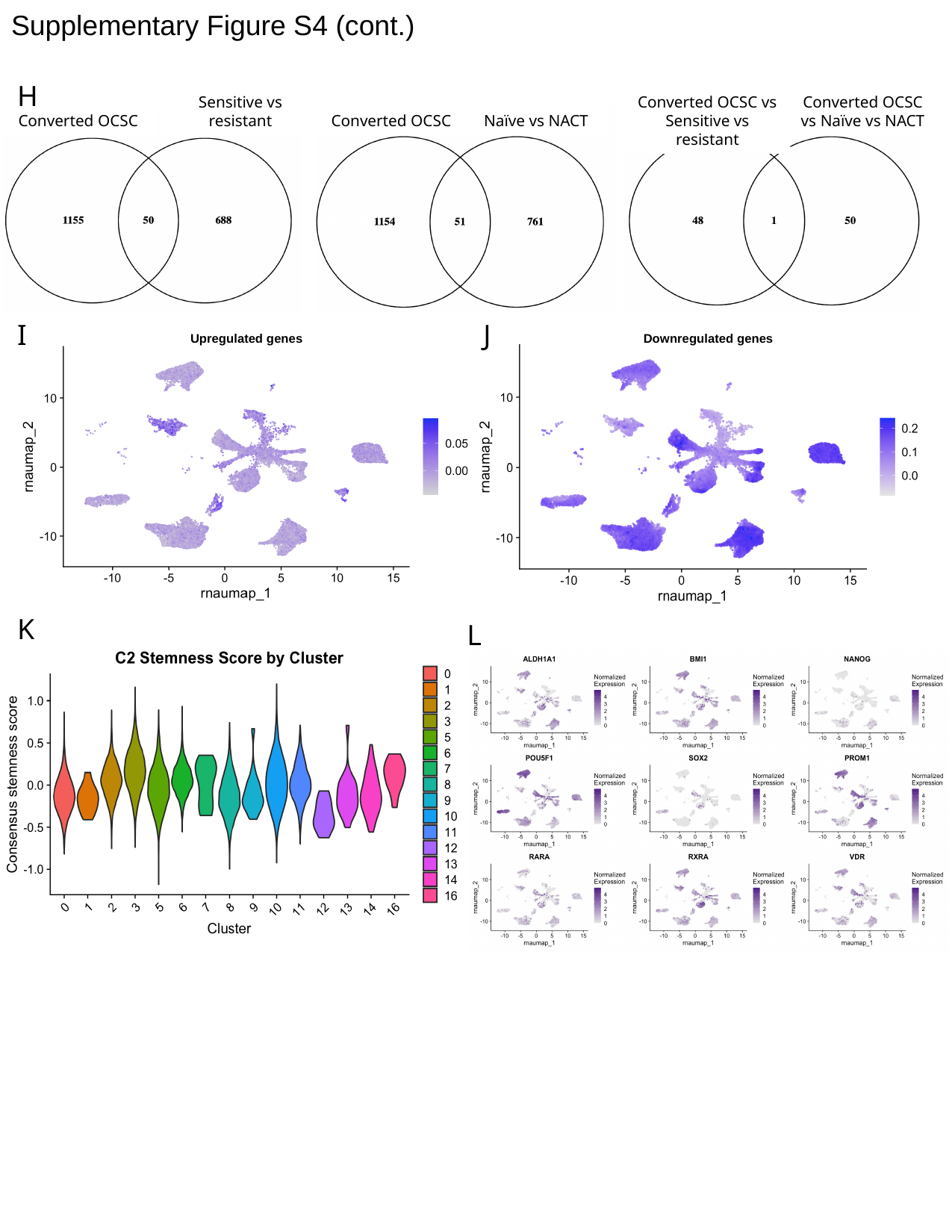

#### Supplementary Figure S4 (cont.)
H
Sensitive vs resistant
Converted OCSC vs Sensitive vs resistant
Converted OCSC vs Naïve vs NACT
Converted OCSC
Converted OCSC
Naïve vs NACT
I
J
Upregulated genes
Downregulated genes
K
L

### Slide 9
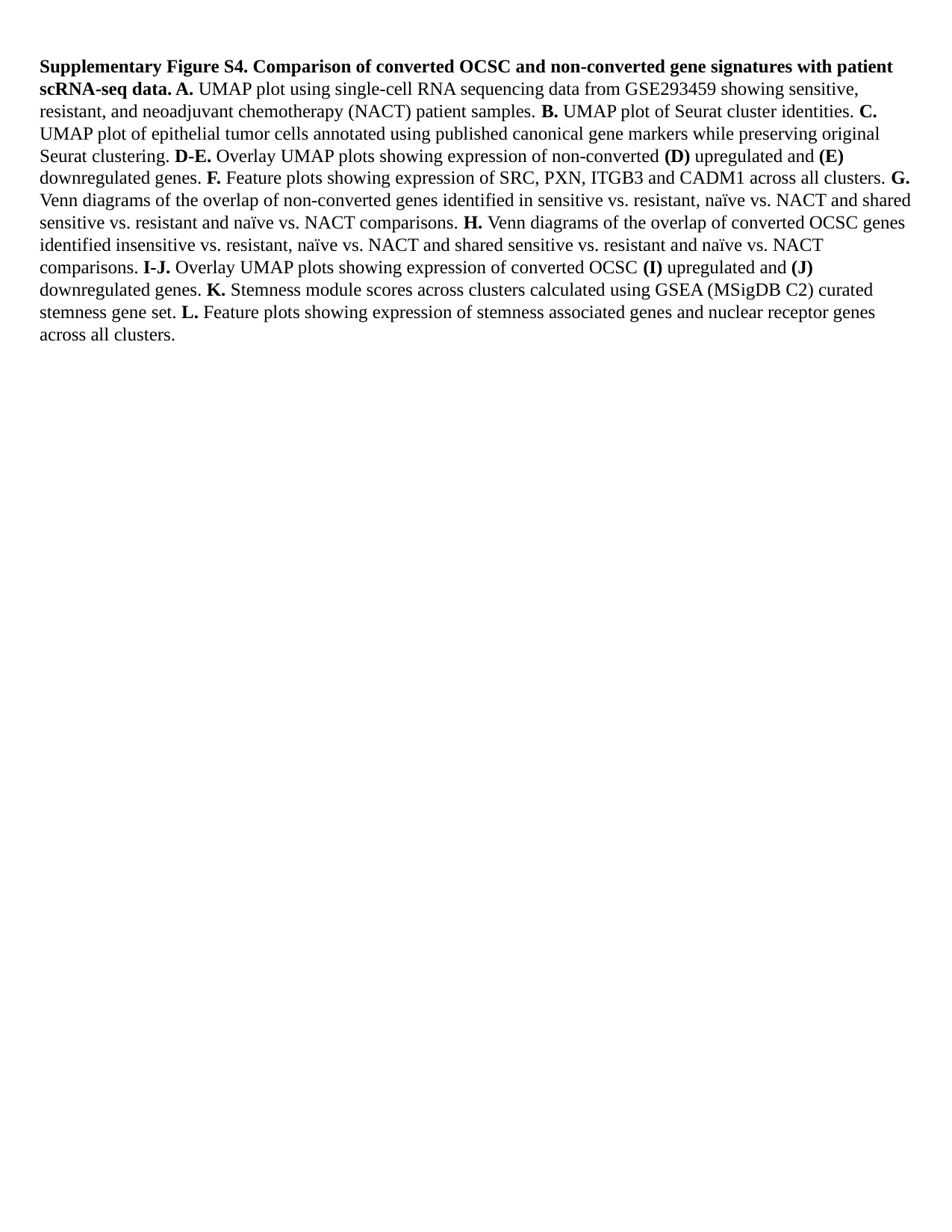

Supplementary Figure S4. Comparison of converted OCSC and non-converted gene signatures with patient scRNA-seq data. A. UMAP plot using single-cell RNA sequencing data from GSE293459 showing sensitive, resistant, and neoadjuvant chemotherapy (NACT) patient samples. B. UMAP plot of Seurat cluster identities. C. UMAP plot of epithelial tumor cells annotated using published canonical gene markers while preserving original Seurat clustering. D-E. Overlay UMAP plots showing expression of non-converted (D) upregulated and (E) downregulated genes. F. Feature plots showing expression of SRC, PXN, ITGB3 and CADM1 across all clusters. G. Venn diagrams of the overlap of non-converted genes identified in sensitive vs. resistant, naïve vs. NACT and shared sensitive vs. resistant and naïve vs. NACT comparisons. H. Venn diagrams of the overlap of converted OCSC genes identified insensitive vs. resistant, naïve vs. NACT and shared sensitive vs. resistant and naïve vs. NACT comparisons. I-J. Overlay UMAP plots showing expression of converted OCSC (I) upregulated and (J) downregulated genes. K. Stemness module scores across clusters calculated using GSEA (MSigDB C2) curated stemness gene set. L. Feature plots showing expression of stemness associated genes and nuclear receptor genes across all clusters.

### Slide 10
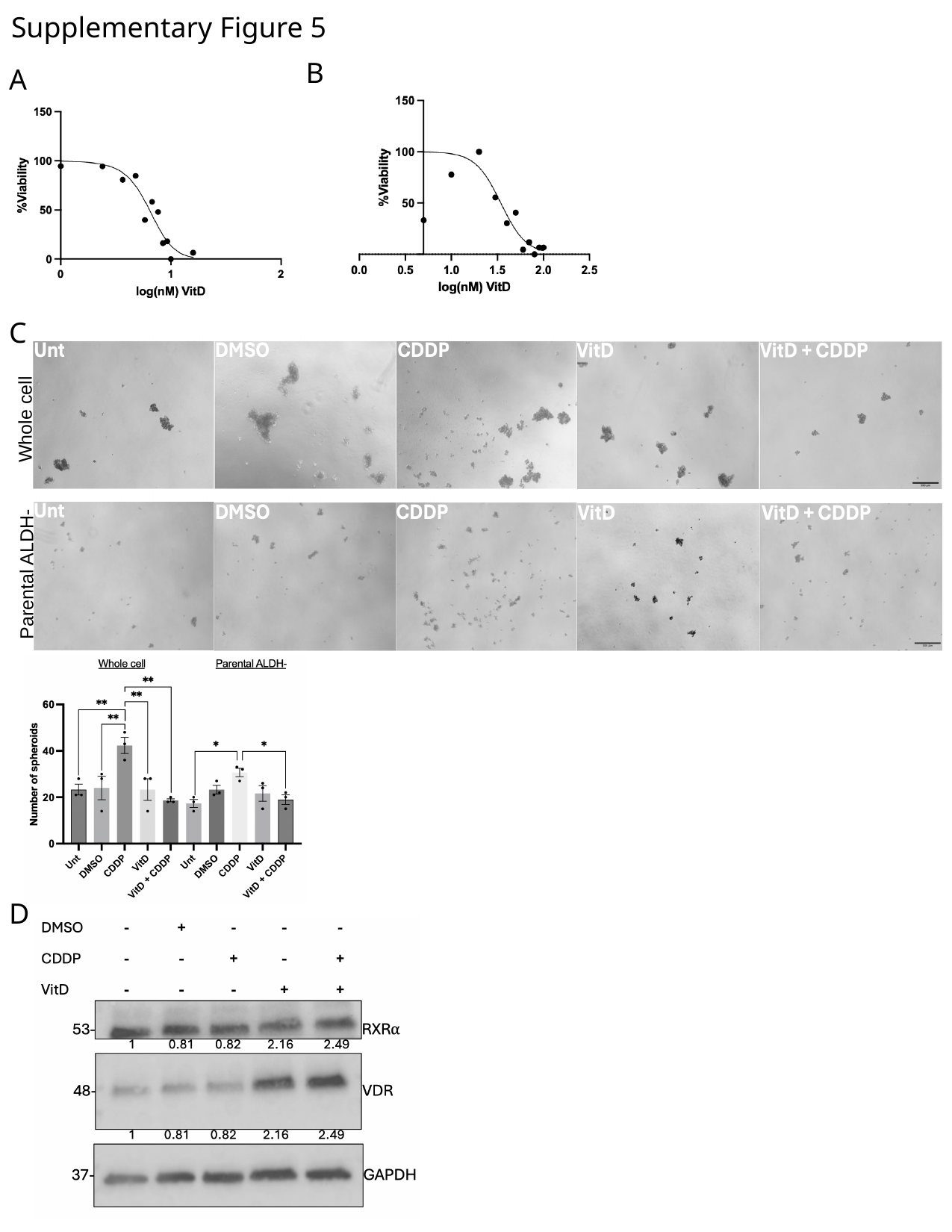

#### Supplementary Figure 5
B
A
C
Whole cell
Parental ALDH-
D

### Slide 11
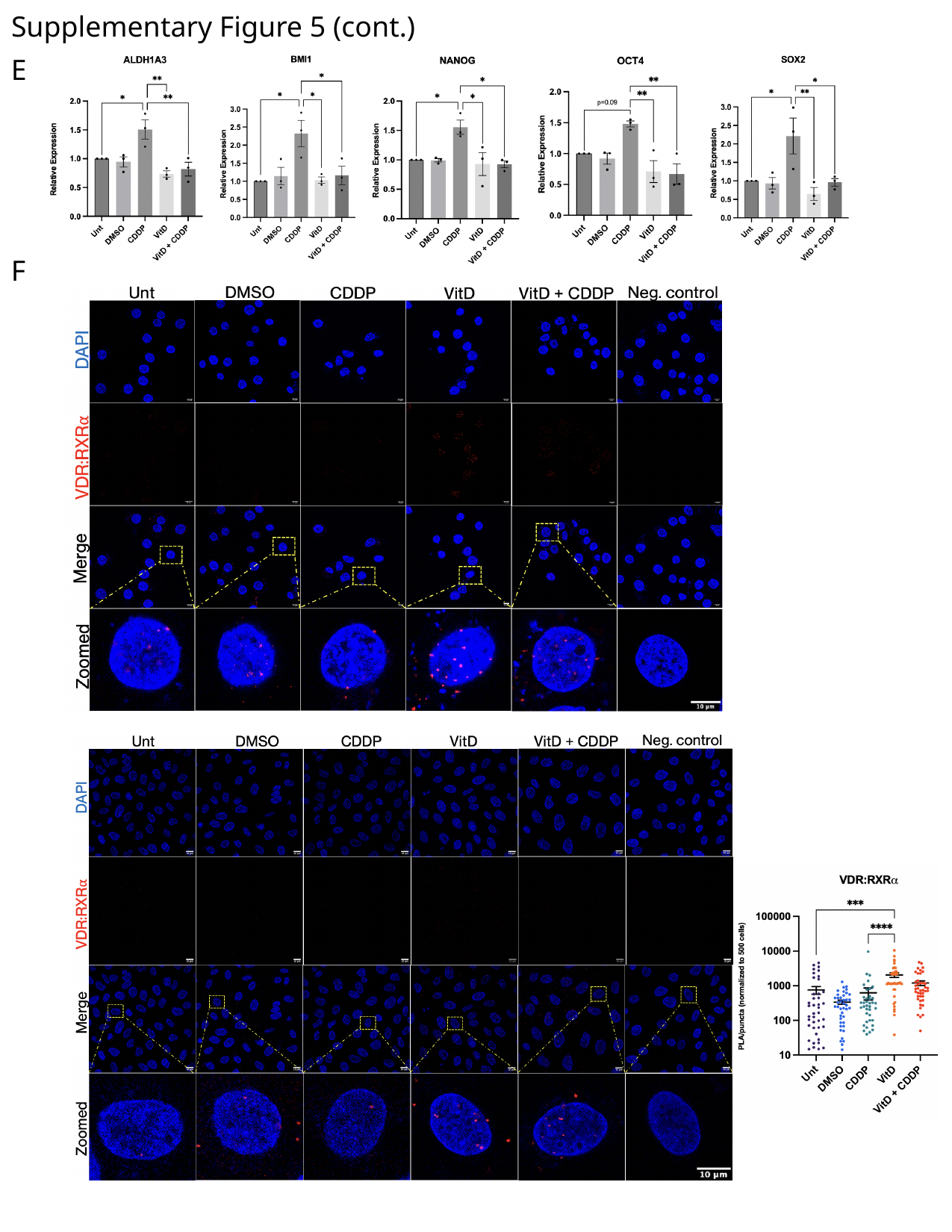

#### Supplementary Figure 5 (cont.)
E
F

### Slide 12
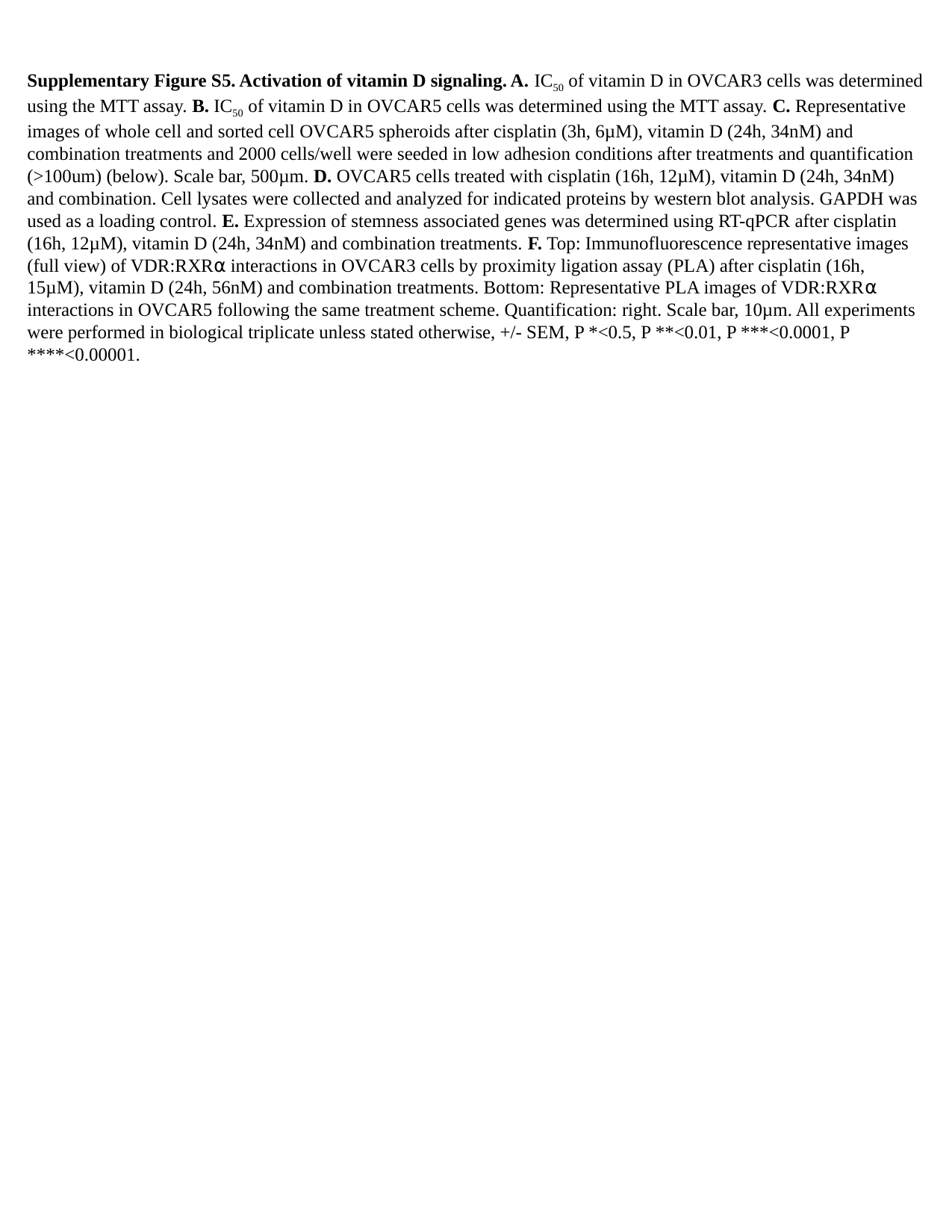

Supplementary Figure S5. Activation of vitamin D signaling. A. IC_50_ of vitamin D in OVCAR3 cells was determined using the MTT assay. B. IC_50_ of vitamin D in OVCAR5 cells was determined using the MTT assay. C. Representative images of whole cell and sorted cell OVCAR5 spheroids after cisplatin (3h, 6µM), vitamin D (24h, 34nM) and combination treatments and 2000 cells/well were seeded in low adhesion conditions after treatments and quantification (>100um) (below). Scale bar, 500µm. D. OVCAR5 cells treated with cisplatin (16h, 12µM), vitamin D (24h, 34nM) and combination. Cell lysates were collected and analyzed for indicated proteins by western blot analysis. GAPDH was used as a loading control. E. Expression of stemness associated genes was determined using RT-qPCR after cisplatin (16h, 12µM), vitamin D (24h, 34nM) and combination treatments. F. Top: Immunofluorescence representative images (full view) of VDR:RXR⍺ interactions in OVCAR3 cells by proximity ligation assay (PLA) after cisplatin (16h, 15µM), vitamin D (24h, 56nM) and combination treatments. Bottom: Representative PLA images of VDR:RXR⍺ interactions in OVCAR5 following the same treatment scheme. Quantification: right. Scale bar, 10µm. All experiments were performed in biological triplicate unless stated otherwise, +/- SEM, P *<0.5, P **<0.01, P ***<0.0001, P ****<0.00001.

### Slide 13
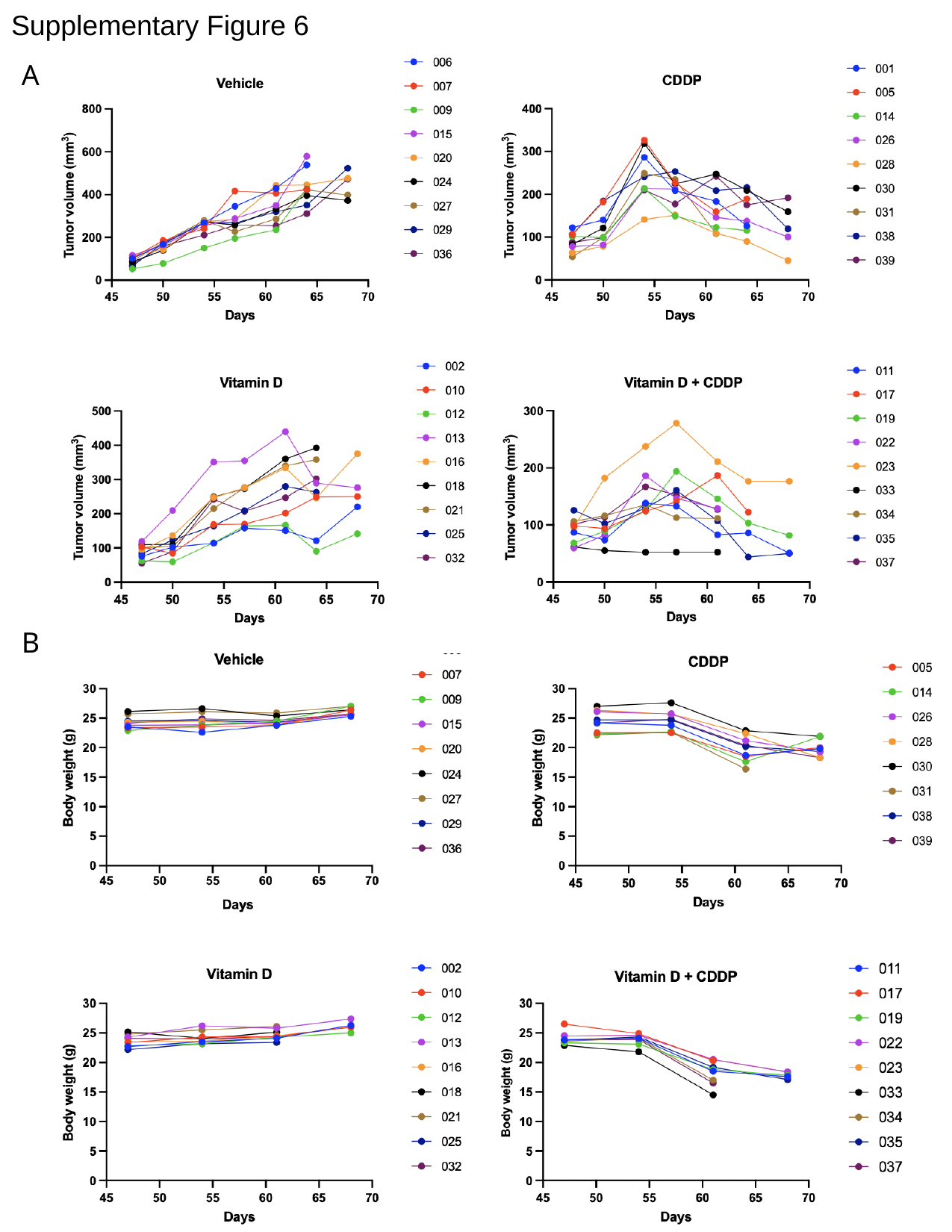

#### Supplementary Figure 6
A
B

### Slide 14
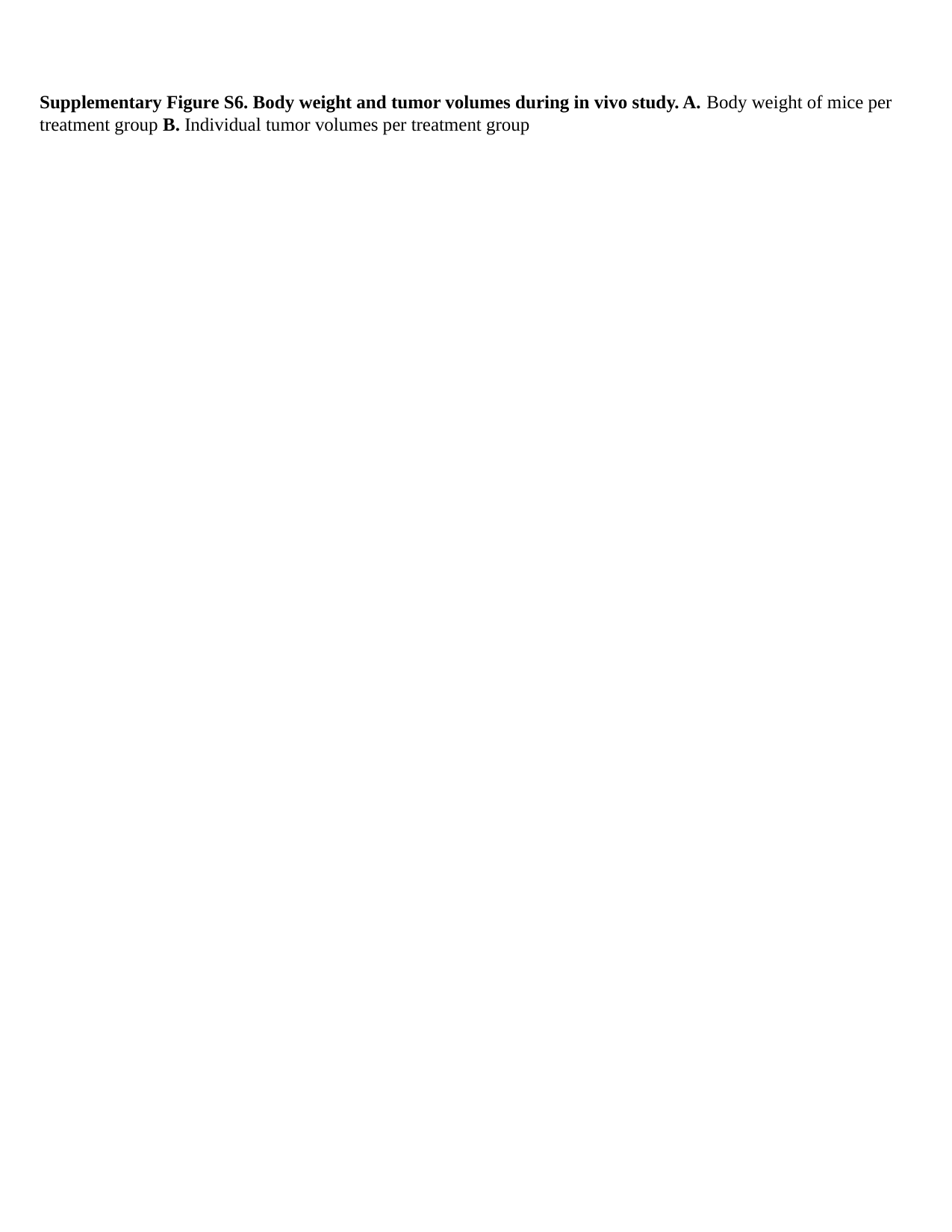

Supplementary Figure S6. Body weight and tumor volumes during in vivo study. A. Body weight of mice per treatment group B. Individual tumor volumes per treatment group

### Slide 15
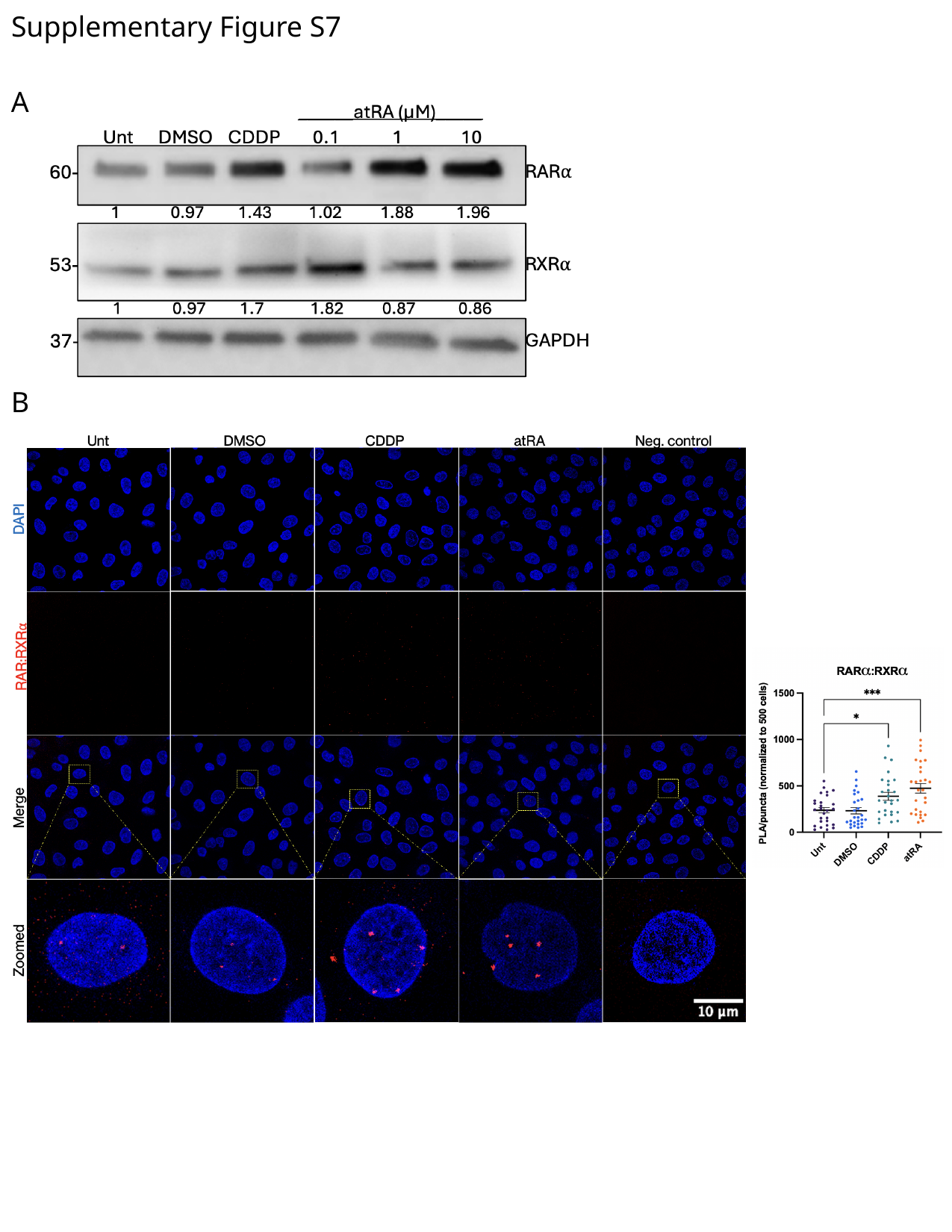

#### Supplementary Figure S7
A
B

### Slide 16
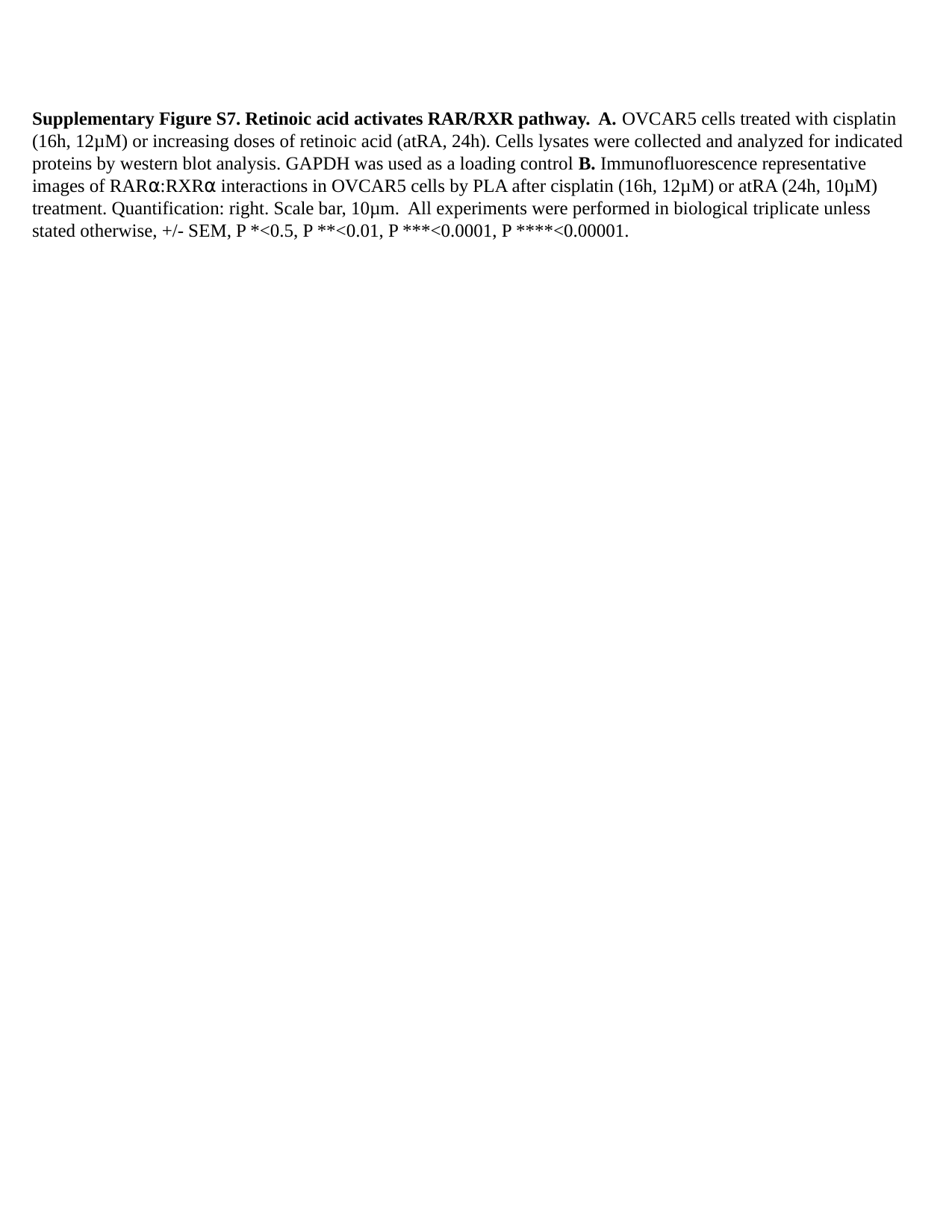

Supplementary Figure S7. Retinoic acid activates RAR/RXR pathway. A. OVCAR5 cells treated with cisplatin (16h, 12µM) or increasing doses of retinoic acid (atRA, 24h). Cells lysates were collected and analyzed for indicated proteins by western blot analysis. GAPDH was used as a loading control B. Immunofluorescence representative images of RAR⍺:RXR⍺ interactions in OVCAR5 cells by PLA after cisplatin (16h, 12µM) or atRA (24h, 10µM) treatment. Quantification: right. Scale bar, 10µm. All experiments were performed in biological triplicate unless stated otherwise, +/- SEM, P *<0.5, P **<0.01, P ***<0.0001, P ****<0.00001.

### Slide 17
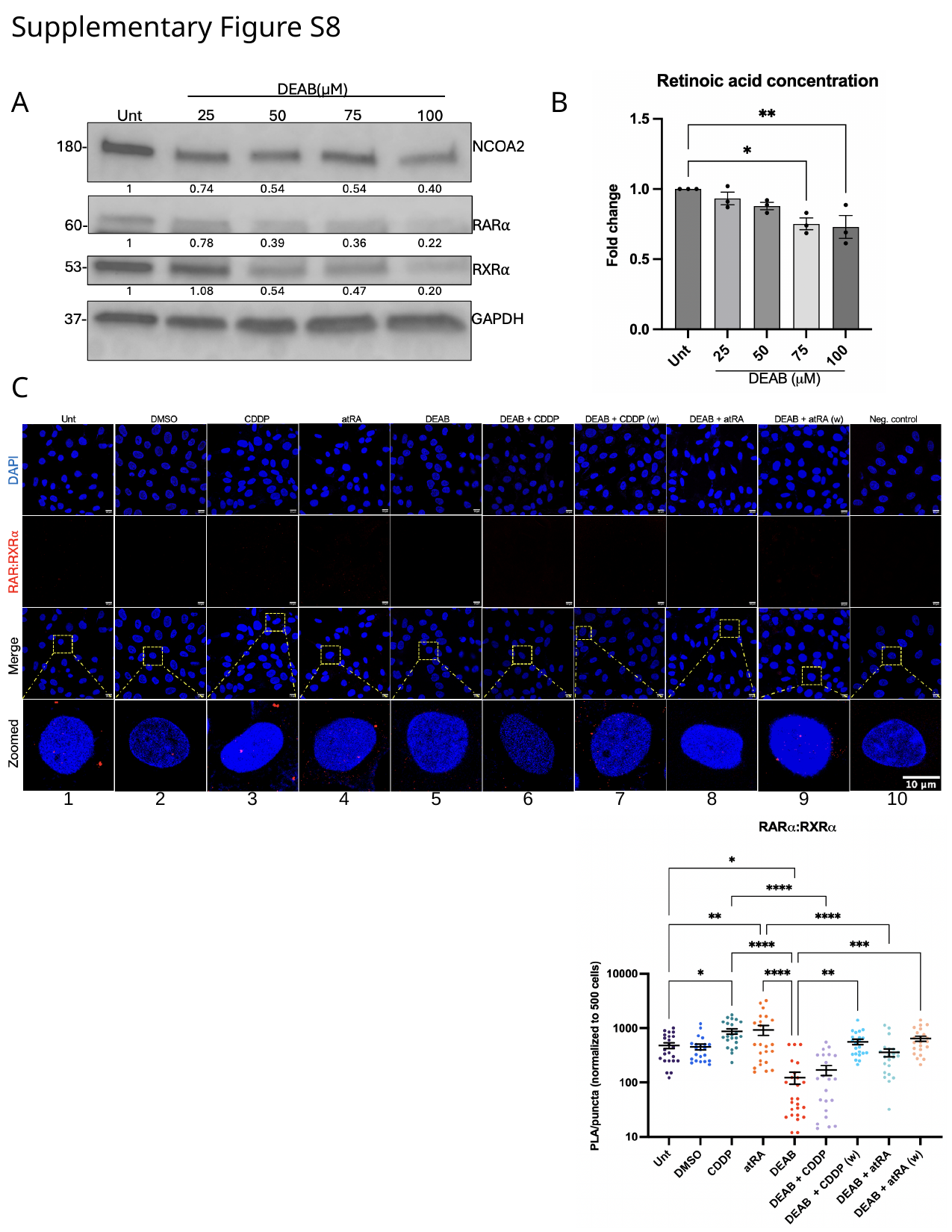

#### Supplementary Figure S8
A
B
C
1
2
3
4
5
6
7
8
9
10

### Slide 18
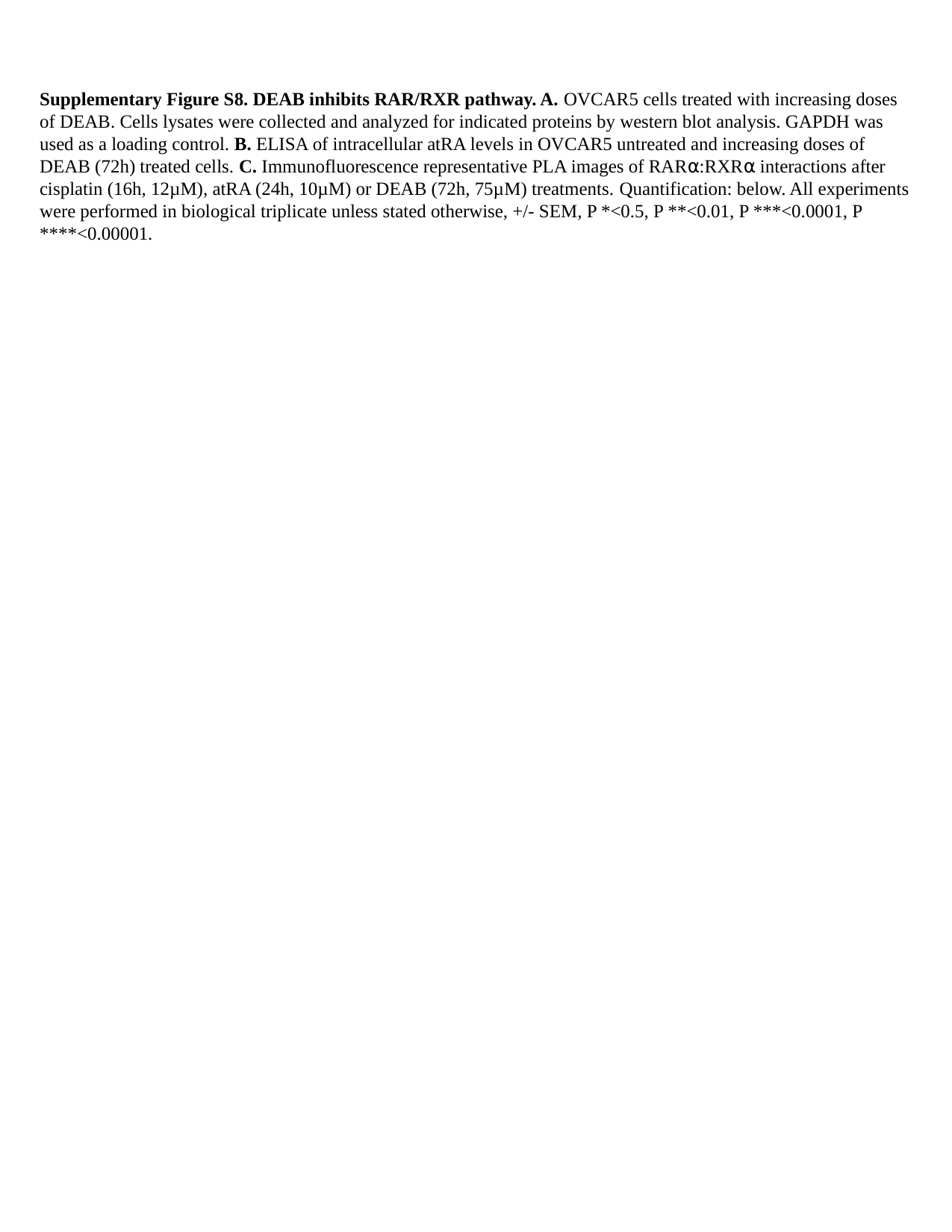

Supplementary Figure S8. DEAB inhibits RAR/RXR pathway. A. OVCAR5 cells treated with increasing doses of DEAB. Cells lysates were collected and analyzed for indicated proteins by western blot analysis. GAPDH was used as a loading control. B. ELISA of intracellular atRA levels in OVCAR5 untreated and increasing doses of DEAB (72h) treated cells. C. Immunofluorescence representative PLA images of RAR⍺:RXR⍺ interactions after cisplatin (16h, 12µM), atRA (24h, 10µM) or DEAB (72h, 75µM) treatments. Quantification: below. All experiments were performed in biological triplicate unless stated otherwise, +/- SEM, P *<0.5, P **<0.01, P ***<0.0001, P ****<0.00001.

### Slide 19
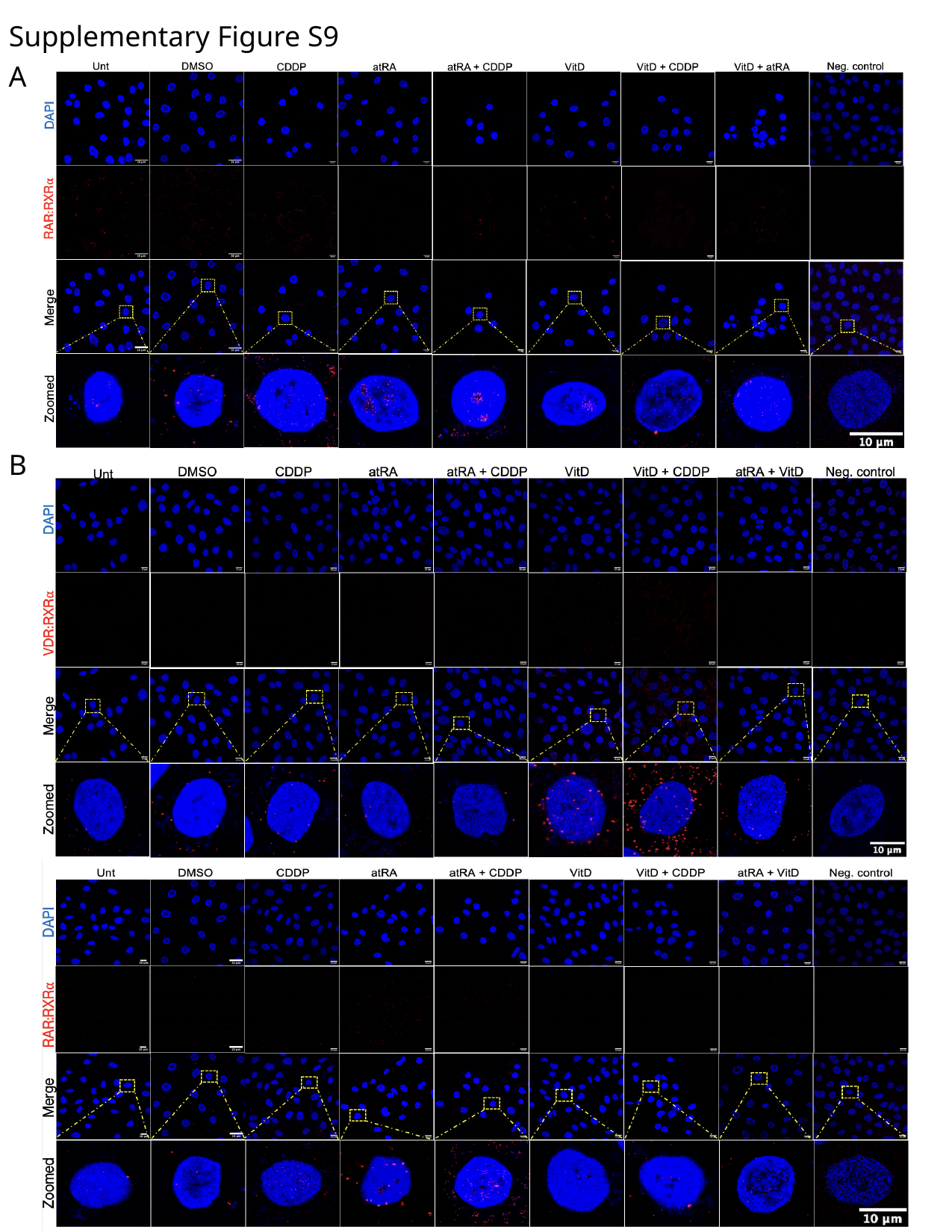

#### Supplementary Figure S9
A
B

### Slide 20
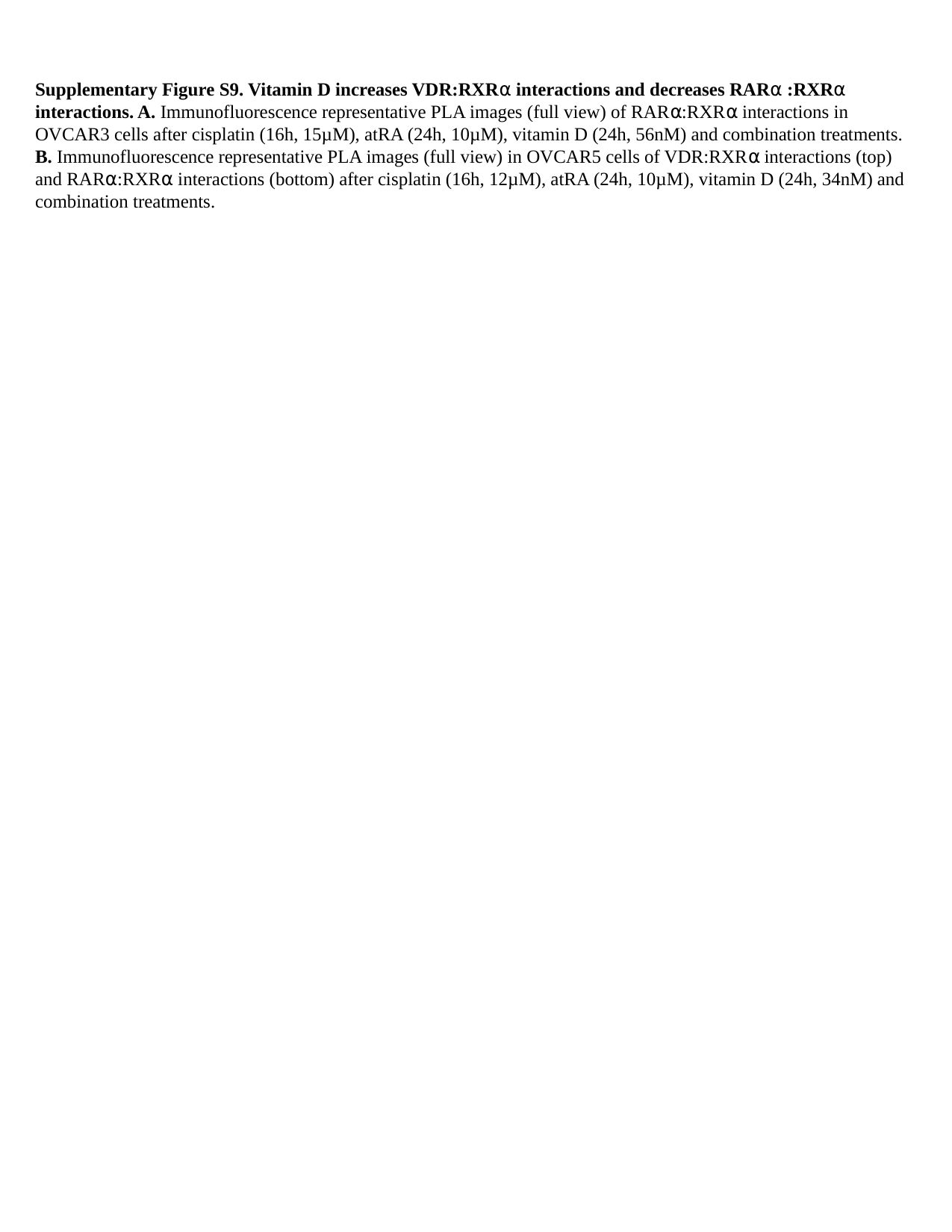

Supplementary Figure S9. Vitamin D increases VDR:RXR⍺ interactions and decreases RAR⍺ :RXR⍺ interactions. A. Immunofluorescence representative PLA images (full view) of RAR⍺:RXR⍺ interactions in OVCAR3 cells after cisplatin (16h, 15µM), atRA (24h, 10µM), vitamin D (24h, 56nM) and combination treatments. B. Immunofluorescence representative PLA images (full view) in OVCAR5 cells of VDR:RXR⍺ interactions (top) and RAR⍺:RXR⍺ interactions (bottom) after cisplatin (16h, 12µM), atRA (24h, 10µM), vitamin D (24h, 34nM) and combination treatments.

### Slide 21
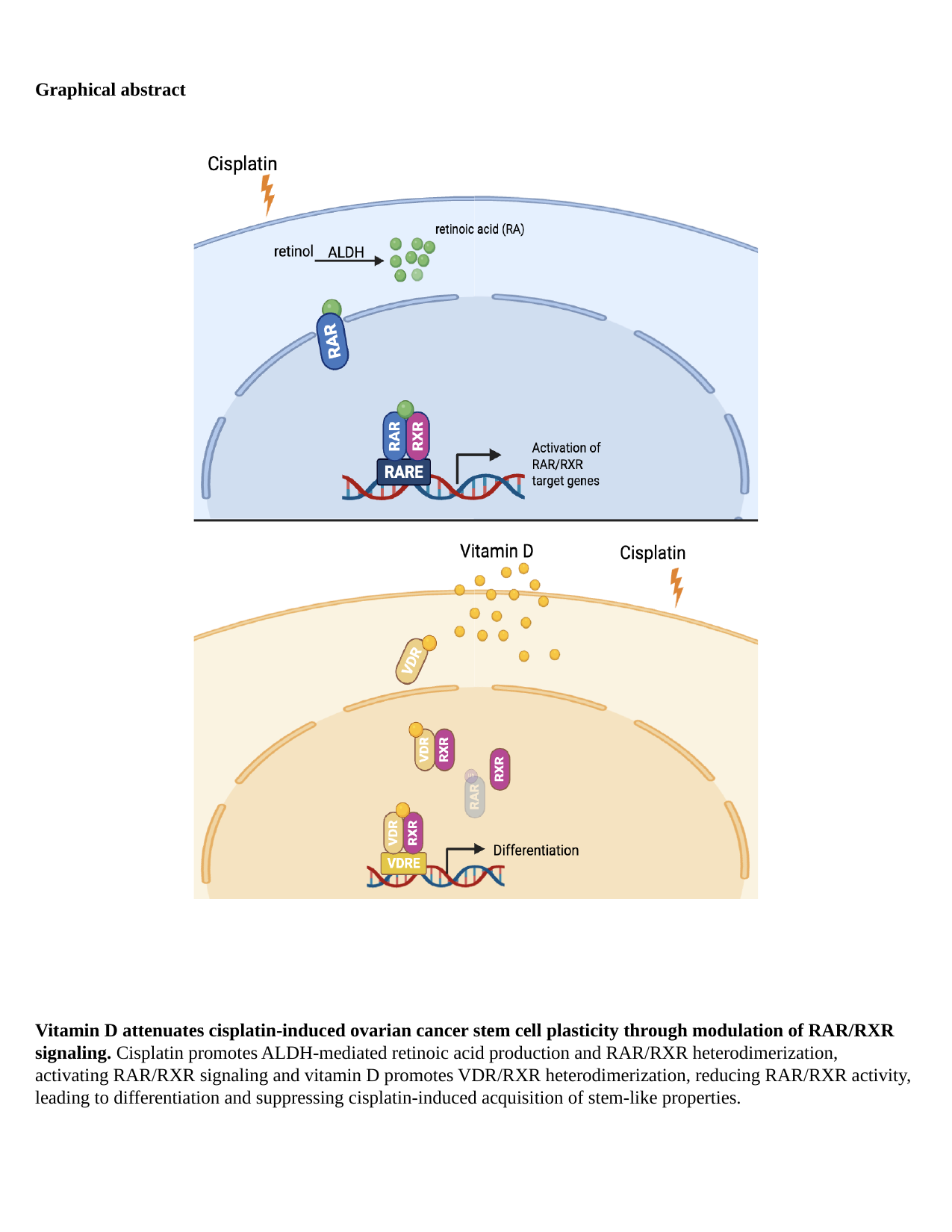

Graphical abstract
Vitamin D attenuates cisplatin-induced ovarian cancer stem cell plasticity through modulation of RAR/RXR signaling. Cisplatin promotes ALDH-mediated retinoic acid production and RAR/RXR heterodimerization, activating RAR/RXR signaling and vitamin D promotes VDR/RXR heterodimerization, reducing RAR/RXR activity, leading to differentiation and suppressing cisplatin-induced acquisition of stem-like properties.
